# Distinct topologically associated domains underlie regulatory logic in lymphatic endothelial cells contributing to proper cell differentiation

**DOI:** 10.1101/2025.06.16.659849

**Authors:** Virginia Panara, Hannah Arnold, Marleen Gloger, Renae Skoczylas, Victoria Vidal Gutierrez, Anna Johansson, Agata Smialowska, Katarzyna Koltowska

## Abstract

The activation and repression of genes is a fundamental part of proper embryonic development and functional tissue formation, ensuring that unique molecular codes are set up to orchestrate cell differentiation. Changes in chromatin organisation dictate accessibility to gene regulatory elements and control gene expression. Several molecular factors regulating lymphatic endothelial cell (LEC) specification and differentiation have been identified. However, it remains to be defined how chromatin is organised in lymphatic endothelium and how it orchestrates lymphatic vessel network formation.

In this study, we combined Hi-C and ATAC-sequencing to characterise 3D chromatin architecture and accessibility in LECs and blood endothelial cells (BECs). We have identified cell type-specific topologically associated domains (TADs) in LECs and BECs. Specifically, our data revealed changes in the TAD boundaries and differentially segregating enhancers regions in lymphatic-associated loci, such as *prox1a* and *tbx1*. This multi-omic approach also defined the regulatory logic for nine genes whose expression is enriched in LECs. *In vivo* validation of their short- and long-range enhancers confirmed their LEC-confined activity. Leveraging these datasets, we reconstructed *mafba* tissue-specific regulatory networks and identified a genetic interaction with *tfe3a in vivo* necessary to limit ectopic vessel formation. Overall, our work provides a powerful resource of multi-omic datasets that can be used to systematically determine the regulatory networks governing LEC identity and genes linked to lymphatic disease.

## Introduction

The lymphatic endothelial cells (LECs) originate from different cellular sources (Hu et al., 2024; Lupu et al., 2025). Despite this, LEC identity acquisition is uniformly accompanied by the induction and regulation of lymphatic gene expression. Changes in chromatin structure and enhancer-promoter interactions are key in controlling gene expression in a cell type-specific manner and serve as an integral regulatory mechanism. However, the mechanisms by which chromatin organisation defines the uniform molecular signature in LECs remain to be understood.

During embryonic development, a pool of lymphatic progenitors gives rise to a lymphatic vascular network that extends across the body. The establishment of this complex and diverse system of vessels depends on steps such as specification, proliferation, migration and differentiation. These processes are regulated by molecular mechanisms which ensure the morphology and functionality of the network. The lymphatic vessel progenitors are distinguished from blood endothelial cells (BECs) by expression of Prox1, a transcription factor (TF) necessary for lymphatic vessel formation in mice and zebrafish (Koltowska et al., 2015a; Wigle et al., 1999). In mice, *Prox1* expression is directly regulated by Sox18 and CoupTFII (François et al., 2008; Srinivasan et al., 2010). Both in mice and zebrafish defined functional lymphatic enhancers contribute to the upstream regulation of *Prox1* (Kazenwadel et al., 2023; Panara et al., 2024).

A key pathway regulating LEC sprouting, specification, proliferation and migration is Vegfc-Vegfr3 signalling. This pathway has an important role in controlling Prox1 expression. In zebrafish, Prox1 is induced by Vegfc-Vegfr3 signalling (Koltowska et al., 2015a), and in mice a Vegfc-Vegfr3 feedback loop regulates the maintenance of Prox1 (Srinivasan et al., 2014). In addition to this role, Vegfc-Vegfr3 signalling also mediates the migration of specified LEC progenitors across the embryo though its downstream components Mafba and Mafbb (Arnold et al., 2022; Koltowska et al., 2015b). Vegfr3 expression itself is regulated by the TFs Tbx1 and Hhex (Chen et al., 2010; Gauvrit et al., 2018). Additional transcription factors, such as Gata2, Foxc2, Foxp2 and Nfatc1, have a described role in the maturation of lymphatic vessels and in lymphatic valve formation (Hernández Vásquez et al., 2021; Kazenwadel et al., 2015; Norrmén et al., 2009; Petrova et al., 2004). Despite the identification of these TFs and signalling pathways as key factors in lymphatic development, the LEC transcriptional network and downstream factors have not been systematically defined.

Central to the regulation of gene expression and the shifts in transcriptomic profiles necessary for cell differentiation and tissue formation are chromatin organisation and enhancer-promoter interactions (Ruiz-Velasco and Zaugg, 2017). In cells, differential gene expression is regulated by chromatin rearrangements that enable DNA-binding proteins, including TF, to access regulatory sequences and interact with gene promoters. Changes in chromatin conformation can occur both at the local level, by the opening or closing of chromatin, and distally, by the formation of chromatin loops connecting distant regions of DNA, such as promoters and their regulatory elements. The loops are further organised in Topologically Associated Domains (TADs) which are restricted by insulators to confine chromatin interactions (Mañes-García et al., 2024). Chromatin architectural changes during lymphatic development have not been explored. Therefore, it is compelling to determine if loop restructuring is one of the molecular mechanisms contributes to the LEC differentiation.

In this study, we have characterised the chromatin organisation of LECs and its associated transcriptional signature using a combination of omics approaches. We uncovered TADs, DNA loops and enhancers specific to the LEC population, providing a more holistic view of the regulatory signature of differentiated LECs. By *in vivo* validation, we have identified active short- and long-range enhancer sequences driving LEC-confined reporter expression. To assess the functional relevance of our data, we have reconstructed the transcriptional network downstream of the known lymphatic TF Mafb. We further validated its genetic interaction with Tfe3a *in vivo*, determining its necessity for regulating ectopic vessel formation. These results show that the acquisition of LEC identity is marked not only by the establishment of a cell type-specific transcriptional signature, but also by a matching rearrangement of chromatin organisation.

## Results

### The LEC and BEC populations present lineage-specific chromatin organisation

To understand how chromatin is organised in lymphatic endothelial cells (LECs) and if it presents unique features compared to blood endothelial cells (BECs), we have used an Hi-C sequencing approach to compare these two cell populations. We isolated LECs and BECs using Fluorescent Activated Cell Sorting (FACS) and sorted cells from *Tg(fli1a:nEGFP)^y7^;TgBAC(prox1a:KalTA4-4xUAS-ADV.E1b:TagRFP)^nim5^*(hereafter referred to as *Tg(prox1a:RFP)^nim5^*) embryos at 5 days post fertilisation (dpf). Double-positive *Tg(fli1a:nEGFP)^y7^*; *Tg(prox1a:RFP)^nim5^* cells were enriched for LECs, while single *Tg(fli1a:nEGFP)^y7^*cells were enriched for BECs, which include both arterial and venous endothelial cell (VEC) populations (**Figure 1A**). Sequencing of 50000 cells per sample from two replicates identified on average 300M number of chromatin contacts per sample. TAD boundaries were called on merged replicates using OnTAD (An et al., 2019), a nested TAD caller. TAD boundaries from all levels in the TAD hierarchy were interrogated for differential contact frequency using TADCompare (Cresswell and Dozmorov, 2020). Consensus TAD boundaries in LEC and BEC, respectively, were identified by comparing contact frequency between replicates (**Table S1-2**). Boundaries not identified as differential between replicates were considered tissue consensus. Consensus TADs were delimited by both consensus boundaries. We found that LECs and BECs have a similar distribution and number of TADs at all hierarchical levels before and after filtering for consensus (**Figure S1A-B**). Similarly, the length distribution of outermost (level 1) TADs before and after filtering shows no marked differences (**Figure 1B-C**), confirming unbiased filtering. We further validated the TADs prediction by TADs length comparison with previously published zebrafish datasets, in dissected brain (Yang et al., 2020) and whole embryo (Franke et al., 2021). In our samples we could observe similar TADs characteristics to those in the dissected tissue data (**Figure S1C**).

**Figure 1:**
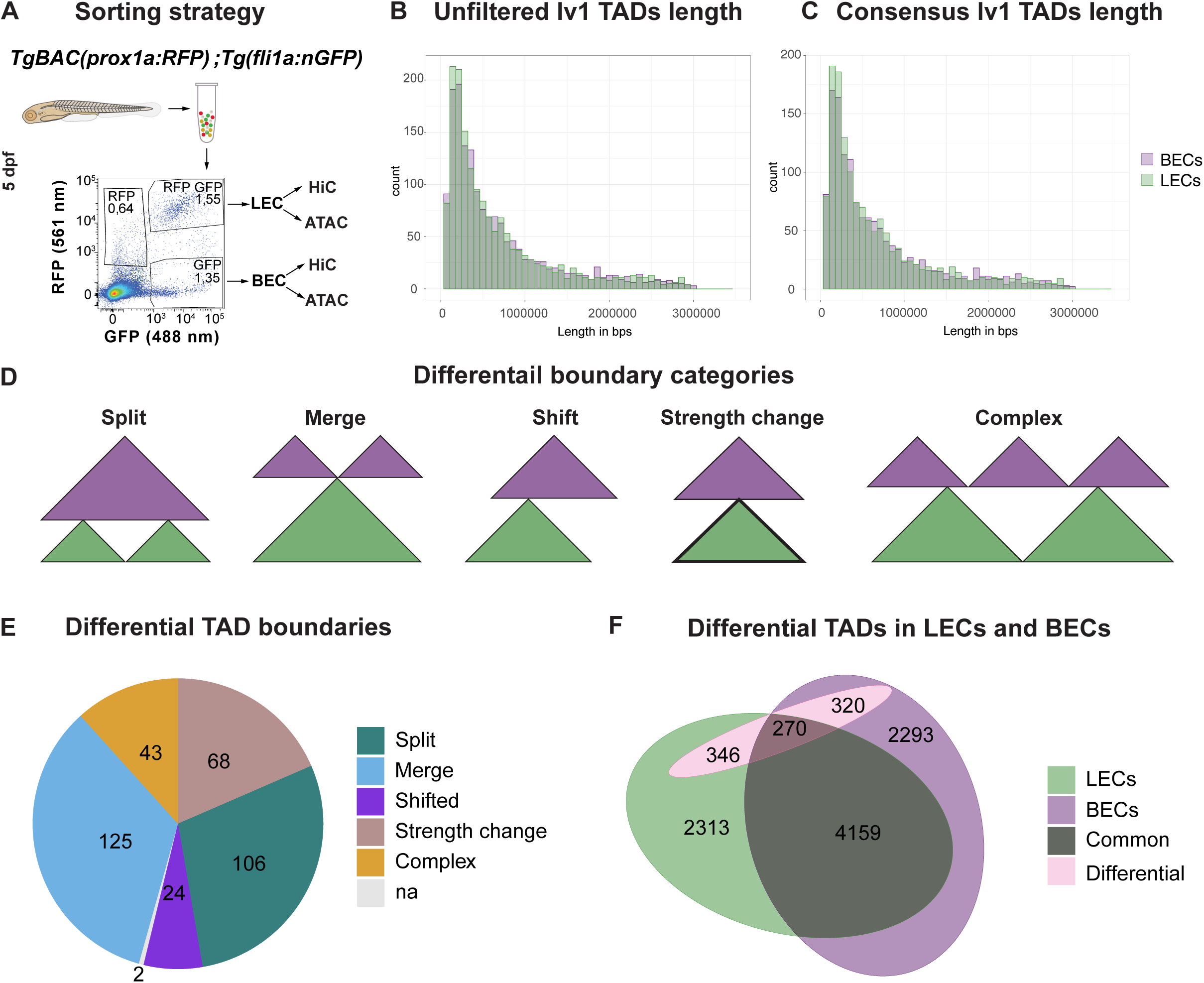
TAD structure presents differences between LECs and BECs. (A) Sorting strategy used for generating the Hi-C and ATAC-sequencing datasets. Three repeats were used for each tissue and analysis. LEC and BEC isolation and sorting strategy from *Tg(fli1a:nEGFP)^y7^; TgBAC(prox1a:KalTA4-4xUAS-ADV.E1b:TagRFP)^nim5^* embryos at 5 days post fertilisation (dpf). (B) Length distribution of unfiltered level 1 TADs identified in LECs and BECs. (C) Length distribution of consensus level 1 TADs identified in LECs and BECs. (D) Representation of the TADcompare categories, adapted from (Cresswell and Dozmorov, 2020). (E) Distribution of the consensus differential boundaries based on the TADcompare categories presented in (D). (F)Summary of consensus differential TADs in LEC and BEC. All lever TADs are considered. Green: Consensus TADs detected in LECs. Purple: Consensus TADs detected in BECs. Pink: Differential TADs.

To characterise the differences in TAD organisation between LECs and BECs, differential TAD boundaries between LECs and BECs were identified by comparing the signal in merged replicate matrices and classified based on published criteria (Cresswell and Dozmorov, 2020) (**Figure 1D**). Consensus TADs delimited by at least one differential boundary in LECs vs. BECs were considered. Overall, 368 differential boundaries were identified between LECs and BECs, of which 43 were classified as “complex” changes, 125 as “merged, 24 as “shift”, 106 as “split” and 24 as “strength” changes” (**Figure 1E**, **Table 3**). To be able to compare TADs between LECs and BECs, we defined common TADs as delimited by both boundaries within 30kb distance in both tissues. In total about 2000 TAD have been identified in LECs and BECs and 4159 common to the two cell populations. Out of these, differential TAD boundaries were found in association with 346 LEC TADs, 320 BEC TADs and 270 common TADs (**Figure 1F**). Overall, these data shows that the global chromatin organisation differs between these two cell types, as shown by the contrasting TAD structure of LECs and BECs.

### The chromatin organisation shows rearrangements in loci of genes enriched in LECs and BECs

To obtain a genome-wide understanding of which LEC-enriched loci undergo chromosomal rearrangements, we have characterised the transcriptional signature of LECs and venous endothelial cells (VECs) by single-cell (sc) RNA-sequencing. We used FACS to isolate a mixed LEC and VEC population by sorting double-positive *Tg(− 5.2lyve1b:Venus)^uu1kk^* and *Tg(fli1a:H2B-mCherry)^uq37bh^* cells at 5 dpf and 7 dpf (**Figure 2A**). To refine the dataset, we used QC filtering and unsupervised clustering (**Figure S2A-B**) which resulted in nine clusters at 5 dpf and seven clusters at 7 dpf (**Figure 2A, S2E**). We identified LEC clusters at 5 and 7 dpf by the expression of known LEC genes, such as *prox1a*, *mafba* and *cdh6* (**Figure 2A, S2C-F, Table S4-5**). The main VEC cluster was determined by the expression of known genes enriched in BECs, such as *cdh5* and *kdrl* (**Figure 2A, S2C-F, Table S4-5**). Further comparisons between LEC and VEC clusters showed enriched expression of *flt4*, *lyve1b* and *tbx1* in the former, whereas expression of *podxl* and *aplnrb* was enriched in the latter. In addition, *fli1a* is expressed in all the clusters, confirming their endothelial identity (**Figure 2A, S2C-F, Table S4-5**). To validate the assigned LEC and VEC clusters, we used transgenic lines reporting the expression of *cdh5* (*TgBAC(ve-cad:ve-cadTS)^uq11bh^*) and *prox1a* (*Tg(prox1a:RFP)^nim5^*) and confirmed that within the endothelium *prox1a* is restricted to lymphatics and *cdh5* to blood endothelium (**Figure 2B**), further validating the identity of the clusters.

**Figure 2:**
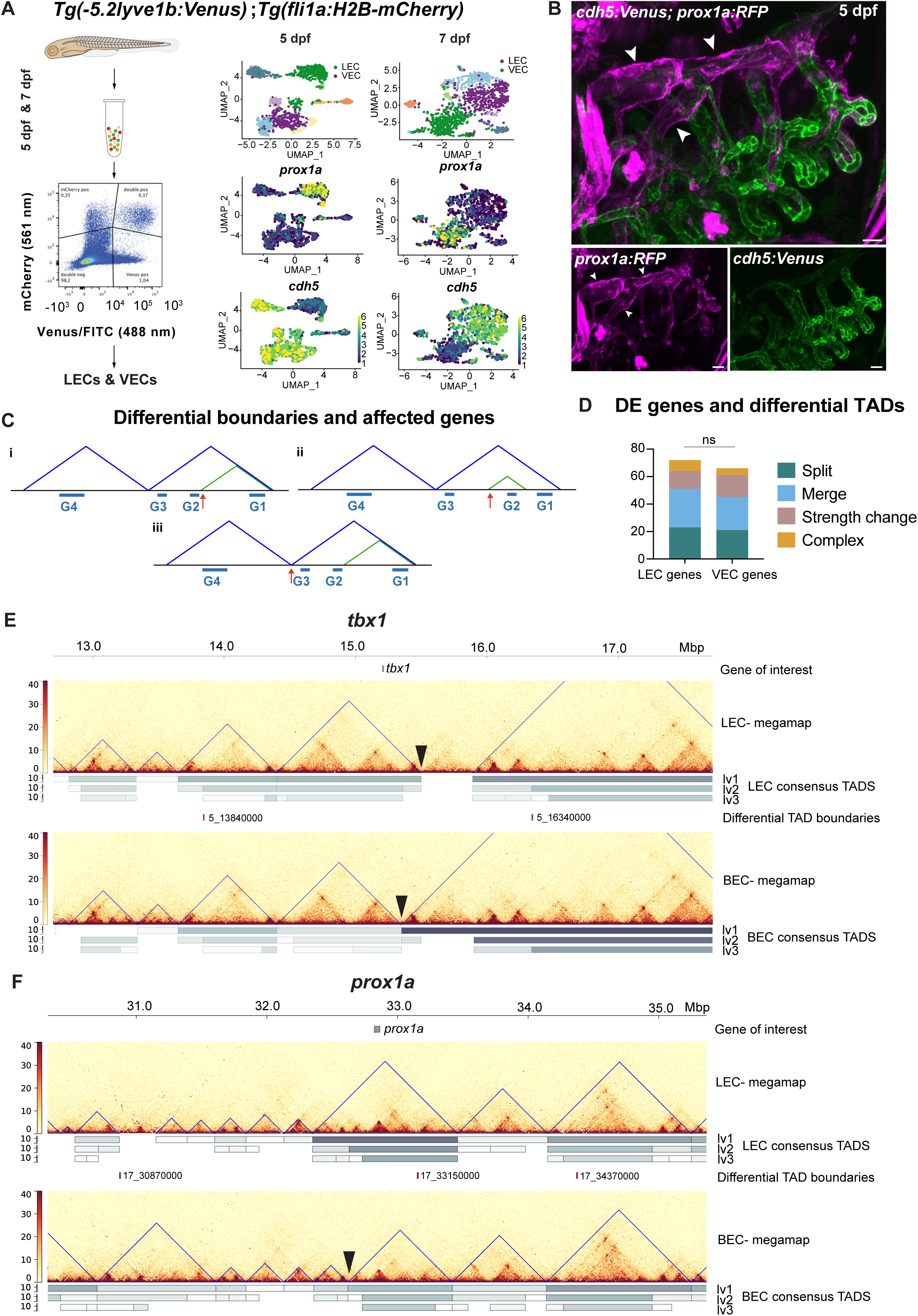
Different chromatin signatures are associated with known lymphatic factors in LECs and BECs. (A) Schematic overview of LEC and VEC isolation and sorting strategy from *Tg(− 5.2lyve1b:Venus)^uu1kk^* and *Tg(fli1a:H2B-mCherry)^uq37bh^*fish for scRNA-sequencing data (top) and identification of the main LEC and VEC clusters with expression plots of known lymphatic marker *prox1a* and known venous marker *cdh5* at 5 dpf and 7 dpf (bottom). Viridis scale colour represents normalised expression. (B) Expression of the transgenic line*s TgBAC*(*ve-cad:ve-cadTS*)*^uq11bh^*(referred to as *cdh5:Venus*) and *TgBAC(prox1a:KalTA4-4xUAS-ADV.E1b:TagRFP)^nim5^ (*referred to as *prox1a:RFP*), showing expression of *chd5* blood endothelium (green) and expression of *prox1a* lymphatic endothelium (magenta, white arrowhead). Scale bars: 20 µm. (C) Schematics of the different gene positions and differential TAD boundaries. Arrow indicates the differential boundary. Triangles colours reflect their different hierarchy level. (D) Distribution of differential TAD boundary in the top 50 DE LEC and BEC genes within level 1 TADs encompassing a differential boundary TAD. (E-F) TAD organisation surrounding the *tbx1* (E) and *prox1a* (F) loci, showing the level 1 unfiltered megaTADs (blue), and the consensus level 1-3 TADs (grey bars) in LECs and BECs. Differential TAD boundaries are plotted in red. Arrowhead: major TAD rearrangements in the proximity of the loci of interest.

We leveraged the scRNA-sequencing data to link the transcriptomic signature of BECs and LECs with the observed changes in chromatin organisation. We quantified the number of genes with LEC- or VEC-biased expression associated with a changing TAD boundary. A gene was considered as LEC or BEC enriched when the |log2FC| > 0.3 and the adjusted p-value < 0.1. Genes potentially affected by a change in TAD boundaries are not only the ones overlapping the boundary itself, but also those located in proximity, as the overall chromatin conformation of the region may be altered. Therefore, this analysis included all outermost TADs (level 1) which are upstream of a lower order differential boundary delimited TAD. Differential level 1 TADs were also included. As an example, in (**Figure 2Ci-ii),** genes G1, G2 and G3 would be considered as loci affected by the changing TAD boundaries, but G4, which is located in a different outermost TAD, would not. On the contrary, in (**Figure 2Ciii)** all genes would be considered affected as the differential boundary is shared by the two outermost TADs. In total, we identified 54 LEC-enriched and 53 BEC-enriched genes associated with changing boundaries (**Figure 2D**).

We then focused on two transcription factors identified by this analysis which have known functions in lymphatic endothelial cells: Tbx1 (Chen et al., 2010) and Prox1a (Koltowska et al., 2015a; Wigle et al., 1999). The *tbx1* locus shows a shift of TAD boundaries, resulting in the inclusion of an additional 150 kbp in the outermost TAD encompassing the gene in LECs (**Figure 2E**). Similarly, the large 1100 kbp outermost TAD encompassing the *prox1a* locus in LECs is split into two smaller TADs in BECs (**Figure 2F**). This indicates that tissue-specific chromatin rearrangements are association with key genes involved in LEC differentiation, and may help to restrict their expression to this cell population.

### Identification of candidate lymphatic regulators based on cell type-enriched gene expression and chromatin accessibility

In addition to large scale chromatin organisation, local changes in chromatin accessibility in LECs and BECs were also characterised. We performed bulk ATAC-sequencing of isolated LECs and BECs from 5 dpf zebrafish, ensuring comparability between the datasets by using the same sorting strategy as for the Hi-C data (**Figure 1A**). We confirmed the high quality of the data by plotting the signal profiles of nucleosome-free and mononucleosome fractions of the data at annotated transcriptional start sites (TSS) (**Figure S3A**). The peak aggregation in relation to TSS showed a standard distribution, and the expected genomic features were detected within the open regions (**Figure S3B),** validating the quality of the dataset. We observed that, out of 86691 peaks, 17908 had reduced accessibility in LECs compared to BECs and 12745 had increased accessibility at FDR cut-off 0.05 (**Figure S3C, Table S6**). Using this data we identified the top 500 differentially accessible peaks (**Figure S3D, Table S7**) and the top 100 promoters (**Figure S3E, Table S8**), showing the presence of differences in the chromatin accessibility between LECs and BECs. To further investigate the relation between chromatin accessibility and transcriptional regulation, we checked if an open chromatin signature in the ATAC-sequencing dataset correlated with a tissue-specific expression enrichment of the closest locus. We found that, for both LECs and BECs, chromatin accessibility and gene expression level are independent from each other (**Figure 3A, Table S7**). Within the top 50 differentially expressed genes (DEG) in LECs and BECs, we found an even split of promoters with increased and decreased accessibility in the same tissue (**Figure S3F, Table S9**). This indicates that the gene expression depends on more than simply chromatin state.

**Figure 3:**
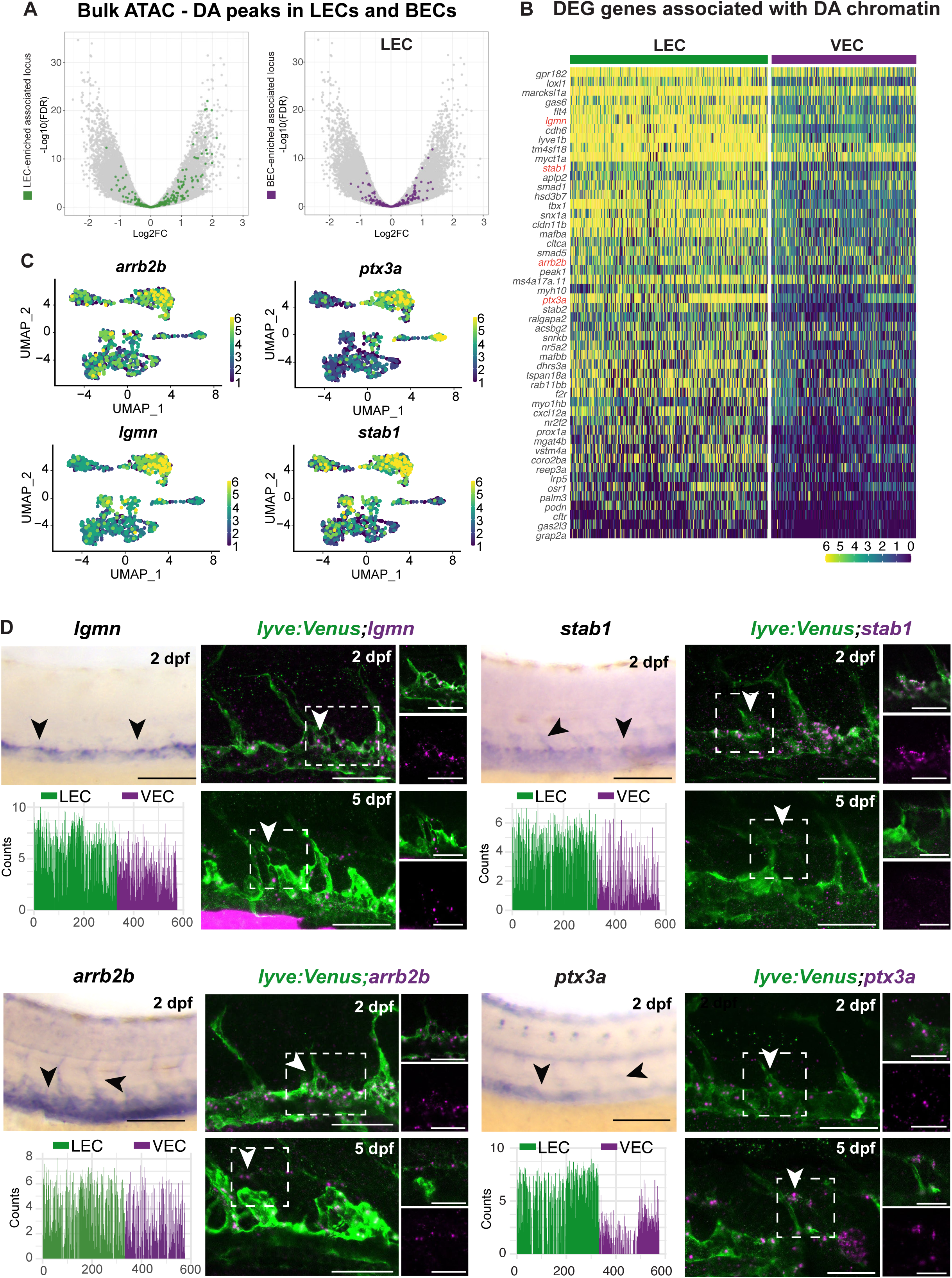
A combination of ATAC-sequencing and scRNA-sequencing data reveals novel LEC factors. (A) Volcano plots of the differentially accessible (DA) peaks in LECs and BECs. Coloured dots indicate the differentially express (DE) status of the nearest locus in the scRNA-sequencing data at 5 dpf. (B) Heatmap showing expression of genes associated with the top DA ATAC-seq peaks at 5 dpf in LECs and VECs. Cells were ordered by their assigned clusters. Colour scale represents log-normalised read counts. (C) Expression of *lgmn*, *stab1, arrb2b* and *ptx3a* at 5 dpf, selected for *in vivo* validation. Vridis scale colour represents normalised expression. (D) Standard and fluorescent *in situ* hybridisation showing endothelial and lymphatic expression of genes from (C) in the trunk at 2 dpf and 5 dpf. Cell counts of gene expression is visualised for the LEC and VEC clusters at 5 dpf. Arrows: endothelial expression at 2 dpf and lymphatic expression at 5 dpf. Boxed area: zoomed panels. Scale bars: 50 µm.

To identify genes associated with both peaks of differentially accessible chromatin and enriched expression in LECs we intersected the ATAC-sequencing and scRNA-sequencing datasets. To do so, we selected the top LEC-specific differentially accessible (DA) regions by average logarithmic fold change (logFC), and annotated them to the name of the closest locus. We then imposed the condition that the genes must be DEG in LECs with log2FC > 0.3 and adjusted p-value < 0.1. We identified 90 genes at 5 dpf and 107 genes at 7 dpf (**Figure 3B, S3G, Table S10-11**) satisfying these criteria. We plotted the top 50 genes by logFC and with an average signal log2CPM > 2 in ATAC-sequencing, ordered by the fraction of cells expressing them in scRNA-sequencing (**Figure 3B, S3G, Table S10-11**). We selected four genes for further characterisation: *arrb2b*, *ptx3a*, *lgmn* and *stab1*. At 5 dpf, their expression is enriched in LECs (**Figure 3C**), and maintained at 7 dpf (**Figure S3H**). To assess in detail the differential chromatin accessibility across the whole locus, we compared the accessibility in three different regions upstream of the transcription start: <= 1kb, 1- 2kb and 2-3kb. We observed that the proximal regions of *ptx3a*, *stab1* and *arrb2b* are more accessible in the BEC population, whereas for all genes the more distal regions (1-2kb and 2-3kb) are more accessible in the LEC population. *lgmn* showed overall more accessibility in LECs (**Figure S3K**). To validate the expression of these genes in the zebrafish lymphatic endothelium we have carried out chromogenic and fluorescent *in situ* hybridisation (**Figure 3D**), and found that all four genes are expressed in the lymphatic endothelium at 2 dpf and 5 dpf in both trunk and face (**Figure 3D, S3I**). Controls with sense probes showed no expression (**Figure S3J**). Hence, by integrating the ATAC-sequencing and scRNA-sequencing datasets, we identified genes presenting both differentially accessible chromatin and enriched expression in LECs, including previously unidentified candidates for lymphatic development, underscoring the robustness and the discovery potential of our datasets.

### LEC chromatin landscape reveals accessible enhancers in the proximity to LEC-enriched genes

Gene expression is regulated through active enhancers marked by an open chromatin conformation. To further validate the ATAC-seq dataset, and to identify lymphatic-specific enhancers, we looked for LEC-specific peaks of open chromatin in the non-coding regions surrounding genes enriched in the LEC clusters (**Figure 4A-B, 4E-F, 4I-J, 4M-N**). We selected four of these putative enhancers to be tested *in vivo* by cloning into the ZED vector (Bessa et al., 2009) (**Figure 4**). Their activity was tested by screening the injected F0 for lymphatic expression. One of the enhancers is located close to *tbx1,* a gene with a well described role in lymphatic development (Chen et al., 2010), while the others are located close to previously undescribed lymphatic candidates: *hyal2b, cxcl12a* and *marcksl1a*. This analysis also identified a previously described (Grimm et al., 2023) *cdh6* lymphatic enhancer (**Figure S4A-C**), validating the potential of the dataset for revealing tissue-specific cis-regulatory sequences. All of the selected enhancers showed expression in the lymphatic endothelium (**Figure 4D, 4H, 4L, 4P**). We further confirmed the identity of the enhancers by verifying that none of the neighbouring loci showed a clear LEC enrichment (**Figure S4E**).

**Figure 4:**
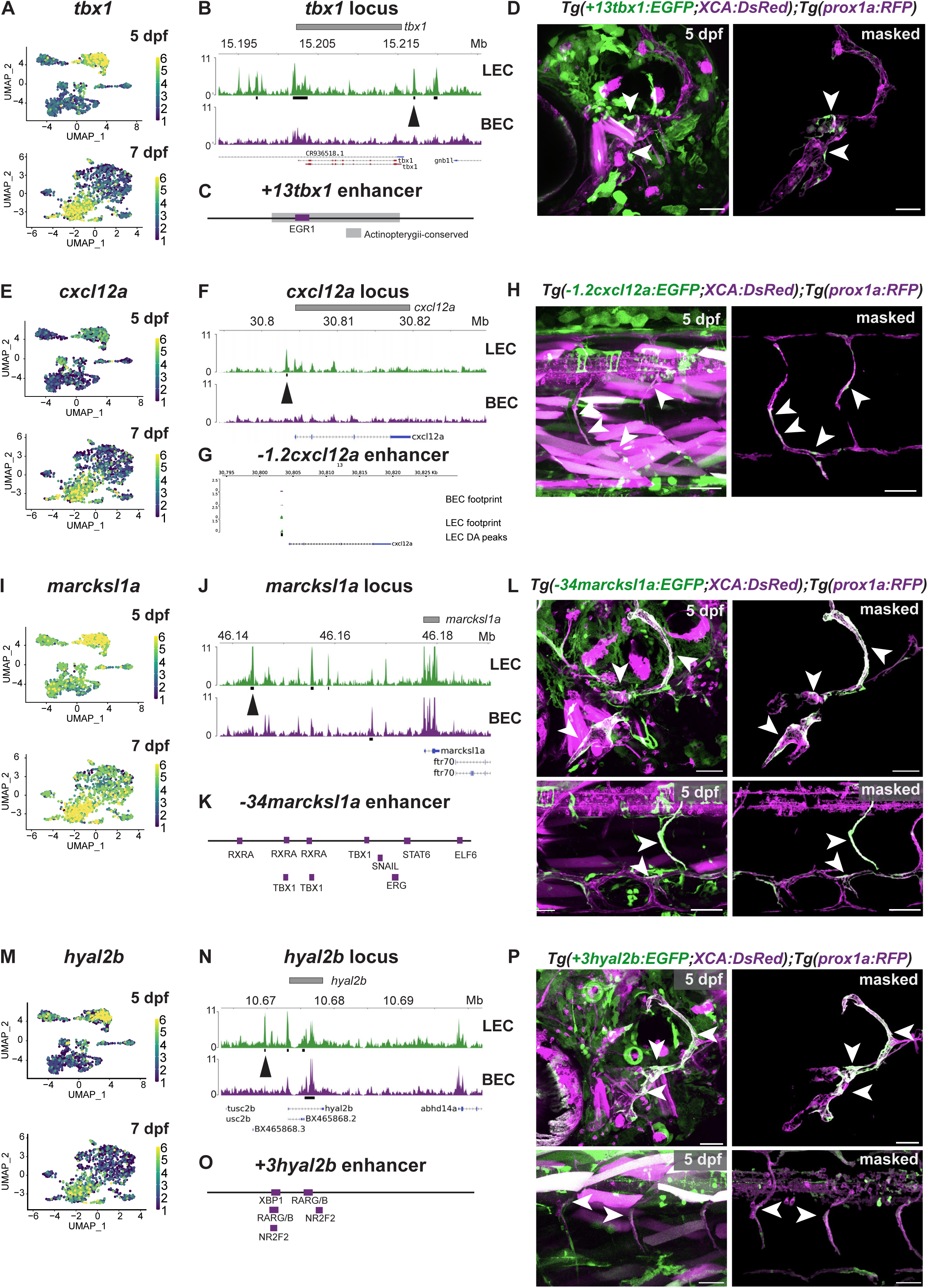
Characterisation of four novel short-range LEC enhancers. (A-D) Characterisation of the *+13tbx1* enhancer. (E-H) Characterisation of the *-1.2cxcl12a* enhancer. (I-L) Characterisation of the *-34marcksl1a* enhancer. (M-P) Characterisation of the *+3hyal2b* enhancer. (A, E, I, M) Enriched expression of the gene in the LEC cluster at 5 dpf and 7 dpf visualised by expression plots. Viridis scale colour represents normalised expression. (B, F, J, N) Chromatin accessibility of the gene locus in LECs and BECs. Black boxes: DA peaks. Arrows: identified enhancers. (C, K; O) Schematics of the identified enhancers. Purple: predicted binding sites of TFs expressed in LECs. Grey: conservation among Actinopterygii. (G): Footprinting signature at the *-1.2cxcl12a* enhancer. (D, H, L, P) Confocal and masked expression driven by the enhancer elements in the lymphatics of 5 dpf embryos. Arrows: expression in the lymphatics. (D) *Tg(+13tbx1:EGFP;XCA:DsRed);Tg(prox1a:RFP)* embryo, showing enhancer activity in the facial lymphatics. (H) *Tg-1.2cxcl12a:EGFP;XCA:DsRed);Tg(prox1a:RFP)* embryo, showing enhancer activity in the trunk lymphatics. (L) *Tg(−34marcksl1a:EGFP;XCA:DsRed);Tg(prox1a:RFP)* embryo, showing enhancer activity in the facial and trunk lymphatics. (P) *Tg(+3hyal2b:EGFP;XCA:DsRed);Tg(prox1a:RFP)* embryo, showing enhancer activity in the facial and trunk lymphatics. Scale bars: 50µm

Some of the tested enhancers, such as *+13tbx1* and *-1.2cxcl12a,* showed activity with in endothelium restricted to a specific lymphatic bed, respectively the facial or the trunk lymphatics (**Figure 4D, 4H**). We assessed the predicted binding of TFs in these enhancers using Transcription factor Occupancy prediction by Investigation of ATAC-sequencing Signal (TOBIAS) and identified a preferential binding site for the endothelial factor EGR1 (Payne et al., 2024) in the *+13tbx1* enhancer (**Figure 4C, S4D**). EGR1 expression is also enriched in LECs at 5 dpf (**Figure S4D**). No predicted TF binding sites were detected using TOBIAS for -*1.2cxcl12a* (**Figure 4G**). On the other hand, -*34marcksl1a* and +3*hyal2b* drove expression in both facial and trunk lymphatic vessels, with additional activity in the blood vascular beds (**Figure 4L-4P**). We identified a range of predicted TF binding sites included *nr2f2, rxra, tbx1, eg1, snail, stat6* and *elf6* for -*34marcksl1a* (**Figure 4K, S4D**) and *xbp1* and *rarga* for +3*hyal2b* (**Figure 4O, S4D**). In conclusion, by using our ATAC-sequencing and scRNA-sequencing datasets, we uncover a set of short-range enhancers that drives the expression in a topologically restricted manner, confined to specific vascular beds.

### The genomic features connected by LEC- and BEC-specific chromatin loops differ

In addition to the identification of TADs, HiC data can be employed to investigate the presence of chromatin loops. These represent strong physical interaction between linearly distant genomic features, and mediate regulatory interactions. Chromatin loops were detected in two independent replicates (**Figure S5A**). The loops present in both replicates within a distance of 25 kb were considered as tissue consensus. Tissue consensus loops without an overlapping or adjacent loop within 40 kb in the other tissue were considered tissue-specific. Overlapping loops and loops within 40 kb in both tissues were considered common loops.

By integrating these lists, we identified the presence of 7061 common loops, 2290 BEC-specific loops and 1095 LEC-specific loops (**Figure S5A, Table S12**). To determine if certain genomic interactions are enriched in the lymphatic endothelium, we characterised the type of features involved in loop formation. First, we identified open chromatin regions placed at the extremities of tissue-specific loops (**Figure S5B**). Quantification of the interactions detected in LEC- and BEC-specific loops determined a total of 1996 interactions specific to LECs and 3462 to BECs (**Figure S5B, Table S13**).

We then characterised the differences in interaction types between LECs and BECs. First, to exclude that the differences in interactions were mirroring underlying differences in the distribution of open chromatin in different genomic features, we used a chi-square test for goodness of fit between observed and expected frequencies. We confirmed that the distribution of genomic features was different from the one expected by randomly picking two open genomic features in the genome (LEC: p<0.0001, BEC: p<0.0001) (**Figure S5C**).

We then used a chi-square test to compare the frequency of different interaction types in LECs and BECs. The analysis confirmed that the distributions were significantly different (p<0.0001), suggesting that not only the location but the type of interactions varies between endothelial cell types. Specifically, the map of the residuals revealed an enrichment of promoter-promoter and promoter-distal interactions in BEC-specific loops, and an enrichment of promoter-intron, promoter-exon, exon-intron, exon-distal, intron-distal and distal-distal interactions in LEC-specific loops (**Figure S5D**). These data revealed distinct differences in the underlying features of loop formation between LECs and BECs, which may shape their divergent developmental trajectories.

### LEC chromatin architecture reveals candidate long-range enhancers interacting with LEC-enriched genes

Next, we wanted to functionally characterise the tissue-specific chromatin loops. Hi-C, ATAC- and scRNA-sequencing data were combined to identify LEC-specific loop-mediated enhancer-promoter interaction. We first defined candidate enhancer sequences as differentially accessible intronic or distal regions of non-coding DNA which interact with the open promoter of a LEC-accessible gene through a LEC-specific loop (**Figure S5E**). The analysis identified 95 loci involved in this kind of interactions. To identify genes whose long-range enhancer interaction coincided with increased expression in the lymphatic endothelium, we intersected the candidate list with genes significantly enriched in LECs in the scRNA-sequencing dataset. This analysis identified 7 genes at 5 dpf and 6 genes at 7dpf (**Figure S5E**). Out of these, *tbx1* (**Figure 5A-B**), *mafba* (**Figure 6B**) and *nr2f2* (**Figure 5E-F**) showed a clear interaction between the promoter and a putative distal regulatory element, and have previously been associated with lymphatic development (Arnold et al., 2022; Chen et al., 2010; Koltowska et al., 2015a; Srinivasan et al., 2010) while *lgmn* (**Figure S6J-O**) has not been previously associated with lymphatic vessels. We then considered whether these long-range interactions and changes in chromatin accessibility are also accompanied by changes in TAD between LECs and BECs, and we found that the loops were associated with TAD boundary shifts at the *tbx1* and *nr2f2* loci (**Figure 5C, 5G**). This change in organisation results in the *+127tbx1* putative enhancer falling outside of the TAD boundary in BECs, whereas in LECs the enhancer and its associated loop are placed within the same TAD as the gene (**Figure 5C**). In the *nr2f2* locus we observed a split resulting in a smaller TAD in BECs, with different loops and differential accessible regions in LECs and BECs (**Figure 5G**).

**Figure 5:**
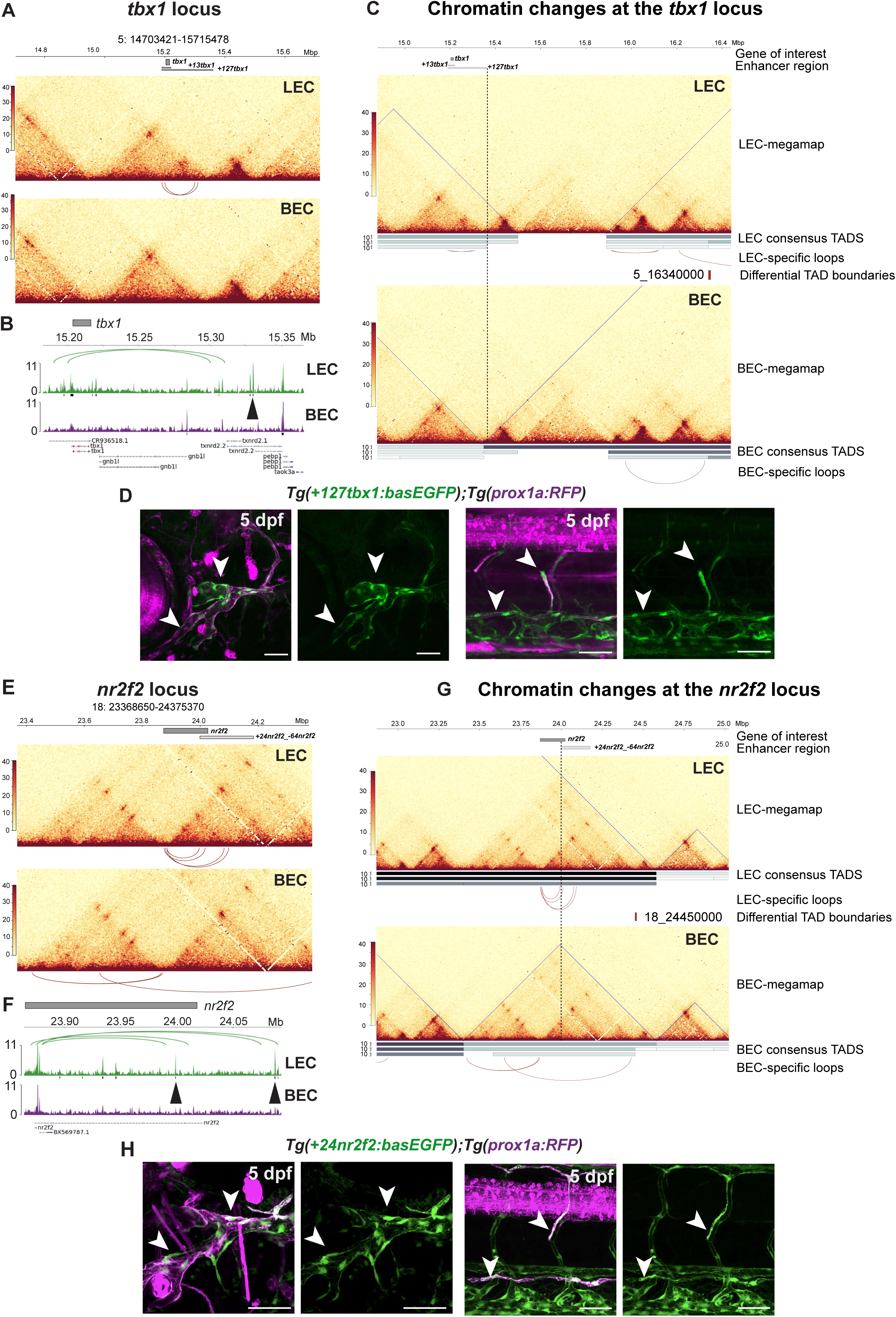
Characterisations of three novel long-range LEC enhancers. (A-D) Characterisations of the *+127tbx1* enhancer. (A) HiC maps in LECs and BECs showing tissue-specific interaction surrounding *tbx1*. Tissue-specific loops are plotted in red. (B) Chromatin accessibility and tissue-specific chromatin loops at *tbx1* in LECs and BECs. Black boxes: DA peaks. Arrow: identified enhancer. (C) Differences in TAD organisation surrounding the *+127tbx1* enhancer in LECs and BECs. Level 1 unfiltered megaTADs shown in blue, consensus level 1-3 TADs shown in grey. Tissue-specific loops are plotted in red. Dotted line indicates the position of the *+127tbx1* enhancer. Differential TAD boundaries are plotted in red. (D) *Tg(+127tbx1:basEGFP);Tg(prox1a:RFP)* confocal and masked expression, showing enhancer activity in the endothelium including the facial and trunk lymphatics at 5 dpf. Arrows: expression in the lymphatics. (E-H) Characterisations of the *+24nr2f2* enhancer. (E) HiC maps in LECs and BECs showing tissue-specific interaction surrounding *nr2f2*. Tissue-specific loops are plotted in red. (F) Chromatin accessibility and tissue-specific chromatin loops at *nr2f2* in LECs and BECs. Black boxes: DA peaks. Arrows: identified enhancers. (G) Differences in TAD organisation surrounding the *+24nr2f2* enhancer in LECs and BECs. Level 1 unfiltered megaTADs shown in blue, consensus level 1-3 TADs shown in grey. Tissue-specific loops are plotted in red. Dotted line indicates the position of *+24nr2f2*. Differential TAD boundaries are plotted in red. (H) *Tg(+24nr2f2:basEGFP);Tg(prox1a:RFP)* confocal and masked expression, showing enhancer activity in endothelium including the facial and trunk lymphatics at 5dpf. Arrows: expression in the lymphatics. Scale bars: 50 µm. (J) Chromatin accessibility and tissue-specific chromatin loops at *lgmn* in LECs and BECs. Black boxes: DA peaks. Arrows: identified enhancer. (K) HiC maps in LECs and BECs showing tissue-specific interaction surrounding *lgmn*. (L) *Tg(+41lgmn:basEGFP);Tg(prox1a:RFP)* confocal and masked expression, showing enhancer activity in the facial lymphatics at 5dpf. Arrows: expression in the lymphatics. (M) Cell counts of the loci surrounding *lgmn*, visualised for LEC and VEC clusters at 5 dpf, showing enriched expression in the selected loci. (N) Schematics of the *+41lgmn* enhancer. Purple: predicted binding sites of TFs expressed in LECs. (O) Heatmap showing the expression in LECs and VECs of the TFs predicted to bind *+41lgmn* at 5 dpf. Cells were ordered by their assigned clusters. Viridis scale colour indicates normalised read counts in log-scale Scale bars: 50µm.

**Figure 6:**
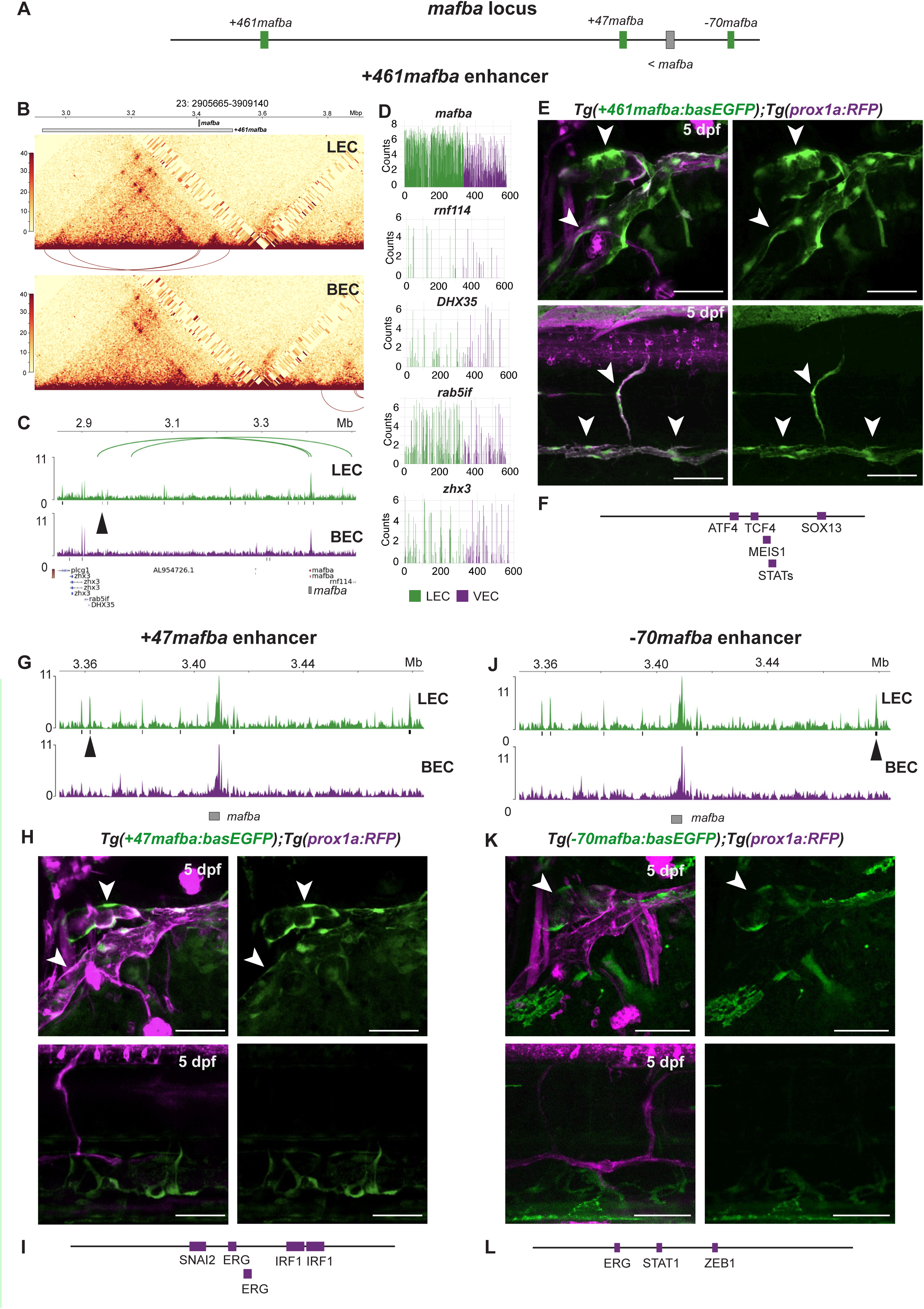
Long- and short-range enhancers of *mafba* drive expression in cell type-specific manner. (A) Schematics of the position of the identified *mafba* enhancers in the surrounding of the *mafba* locus. (B-F) Characterisations of the *+461mafba* enhancer. (B) HiC maps in LECs and BECs showing tissue-specific interaction surrounding *mafba*. Tissue-specific loops are plotted in red. (C) Chromatin accessibility and tissue-specific chromatin loops at the *mafba* locus in LECs and BECs. Black boxes: DA peaks. Arrows: *+461mafba* enhancer. (D) Cell counts of the loci close to *+461mafba*, visualised for LEC and VEC clusters at 5 dpf, showing enriched LEC expression of *mafba*. (E) *Tg(+461mafba:basEGFP);Tg(prox1a:RFP)* embryo, showing enhancer activity in the facial and trunk lymphatics. (F) Schematics of the *+461mafba* enhancer. Purple: predicted binding sites of TFs expressed in LECs. (G-I) Characterisations of the *+47mafba* enhancer. (G) Chromatin accessibility at the *mafba* locus in LECs and BECs. Black boxes: DA peaks. Arrow: *+47mafba* enhancer. (H) *Tg(+47mafba:basEGFP);Tg(prox1a:RFP)* embryo at 5 dpf, showing enhancer activity in the facial lymphatics. (I) Schematics of the *+47mafba* enhancer. Purple: predicted binding sites of TFs expressed in LECs. Arrows: expression in the lymphatics (J-L) Characterisation of the *-70mafba* enhancer. (J) Chromatin accessibility at the *mafba* locus in LECs and BECs. Black boxes: DA peaks. Arrow: *-70mafba* enhancer. (K) *Tg(−70mafba:basEGFP);Tg(prox1a:RFP)* embryo at 5 dpf, showing enhancer activity in the facial lymphatics. Arrows: expression in the lymphatics (L) Schematics of the *-70mafba* enhancer. Purple: predicted binding sites of TFs expressed in LECs Scale bars: 50 µm.

To validate the long-range enhancers *in vivo*, we generated constructs using a vector with an *e1b TATA* minimal promoter (Villefranc et al., 2007), as we have previously confirmed that the empty plasmid does not drive any vascular expression (Panara et al., 2024). An analysis of the stable line for *+127tbx1* showed expression in the facial and trunk lymphatic vessels, as well as the posterior cardinal vein (**Figure 5D**). As this enhancer sits 127 kb downstream of *tbx1,* we checked the expression of the surrounding genes in endothelial cells using the scRNA-sequencing dataset, founding minimal expression of *txtrd2.1, txtrd2.2* and *pebp1* in both LECs and VECs, further supporting the specificity of the enhancer-driven expression (**Figure S6A**). An mVista analysis revealed a restricted evolutionary conservation of this element, which is found only within Actinopterygii (**Figure S6B**). The analysis of predicted TF binding sites showed the presence of sites for ERG, TFC/L2 and SOX family members, all of which are expressed in the LEC and VEC clusters (**Figure S6B-C**).

For *nr2f2* we identified two enhances: one located 24 kb downstream and one 64 kb upstream from the transcription start site (**Figure 5E-F**). The *+24nr2f2* enhancer drove expression in facial lymphatics, as well as in arterial and venous vasculature (**Figure 5H**). The *-64nr2f2* enhancer showed no expression in the endothelium (**Figure S6G**). The surrounding genes (*lysmd4* and *nctp2a*) were not expressed in the LEC and VEC clusters in the 5 dpf scRNA-sequencing dataset (**Figure S6D**). The predicted binding sites of TFs expressed by LECs included NR1D2, NR5A2, ERG and RPBJ for *+24nf2f2* and ERG, KLF2, KLF7, ELK3, EGR1, FEV and HOXD9 for *-64nr2f2* (**Figure S6E, S6H**). Both of the enhancers were conserved in Actinopterygii in the mVISTA analysis (**Figure S6F, S6I**).

For *lgmn* we have identified a long-range enhancer 41 kb downstream of the locus (**Figure S6J-K**) driving expression only in the facial lymphatics but not in the trunk (**Figure S6L**). The surrounding genes (*zgc:153911, pcnx1, si:dkey-157l19.2*) did not show expression in the LEC and VEC transcriptomic data at 5 dpf (**Figure S6M**). The predicted TF binding sites included members of the SOX and retinoic acid family as well as ERG and RPBJ (**Figure S6N**). All of these genes are expressed in the LEC clusters (**Figure S6O**) and have been implicated in lymphatic or vascular formation, further supporting their relevance to these processes.

By leveraging chromatin looping we identified long-range enhancers that drive tissue-specific expression in the endothelium, revealing a previously unappreciated multifaceted nature of transcriptional regulation of lymphatic-enriched genes. Notably, two of these genes are located within regions subject to differential TAD formation in LECs and BECs, implicating different chromatin conformation as a mechanism for regulating cell lineage restricted expression. Together these data reveal a complex regulatory landscape in which lymphatic-enriched gene expression is orchestrated by chromatin organisation through local enhancers, long-range interactions, or a combination of both.

### Long- and short-range enhancers of mafba drive expression in a cell type- specific manner

To determine how the intricate regulatory landscape is linked to the regulation of key transcription factors for specific cellular processes, we focused on *mafba*. *mafba* expression has been reported in both LECs and VECs by *in situ* hybridisation and is necessary for LEC migration (Koltowska et al., 2015b). The scRNA-sequencing revealed that *mafba* is differentially expressed between LECs and VECs, with higher expression in LECs at both 5 and 7 dpf (**Figure S2D-S2F**). To understand how this gene regulation is achieved in LECs and VECs, we have focused on identifying its vascular enhancers. Although no TAD rearrangements were detected in the *mafba* locus, we identified three candidate sequences, *+461mafba*, *+47mafba* and *-70mafba* (**Figure 6A-C, 6G, 6J**), based on a combination of ATAC-sequencing and Hi-C. Tested *in vivo*, all three candidates can drive the expression in the facial lymphatic vessels, with *-70mafba* being more restricted to the facial collecting lymphatic vessel (**Figure 6E, 6H, 6K**). Expression in the trunk lymphatic vessels was observed for *+461mafba* (**Figure 6E**), whereas blood endothelial expression was observed for *+47mafba* and *- 70mafba* (**Figure 6H, 6K**). The predicted binding sites for TFs expressed in LECs included ATF4, TCF4, MEIS1, SOX13 and STATs for *+461mafba* (**Figure 6F, S7A**); SNAI2, ERG and IRF1 for *+47mafba* (**Figure 6I, S7A**), and ERG, STAT1 and ZEB1 for *-70mafba* (**Figure 6L, S7A**). Among these, ATF4, STATs and SNAI2 have not been implicated in regulation of lymphatic vessel formation. Interestingly, the *+461mafba* enhancer falls at the end of a LEC-specific loop, supporting direct chromatin interaction with the *mafba* promoter (**Figure 6B-C**). However, this element is placed 461 kilobases away from *mafba* and close to other genes. Therefore, we checked the expression of these loci and found only a small percentage of LECs expressing *rnf114*, *DHX35, rebi5if, zhx3 and plcg1*. When compared with the robust expression of *mafba* (**Figure 6D**), this provides further evidence of the enhancer’s role as a regulator of *mafba* in LECs.

Together, these data highlight the importance of considering chromatin loops in the reconstruction of gene regulation, as spatially distant enhancers may work in concert to fine-tune transcriptional control. The complexity of *mafba* regulation, with multiple enhancers being capable of driving the expression in different vascular beds, suggests that coordinated enhancer activity may be critical for modulating lineage-specific expression. This complex regulation of *mafba, a* key factor for LEC migration, further emphasises the intricate link between enhancer activity and cellular functions.

### Specific predicted TF binding sites are enriched in LECs and BECs at 5 and 7 dpf

Building on our identification of tissue-specific transcriptional signatures and chromatin organisation, we next explored the regulatory networks that establish and maintain the gene expression programs of LECs and BECs using our multi-omic datasets. To do so, we first identified the molecular factors which may bind to the accessible chromatin regions. We analysed the TF footprints in promoter ATAC peaks to predict the TF binding sites preferentially occupied in each cell type using the TOBIAS tool (**Figure 7A, Table S14**). A proportion of the genes associated with these promoters showed upregulated expression in LECs or BECs (**Figure 7A, Table S14**). We then asked whether the TFs whose motifs were preferentially occupied in LECs and BECs were also enriched in their corresponding cell-type in the scRNA-sequencing dataset. However, with the exception of *nr2f2* (which is both preferentially occupied and enriched in LECs), all of the TFs whose motifs were preferentially occupied in one tissue showed comparable expression levels in LECs and VECs (**Figure 7B, Table S15-16)**, suggesting that TF RNA levels alone do not necessarily reflect their biological relevance.

**Figure 7:**
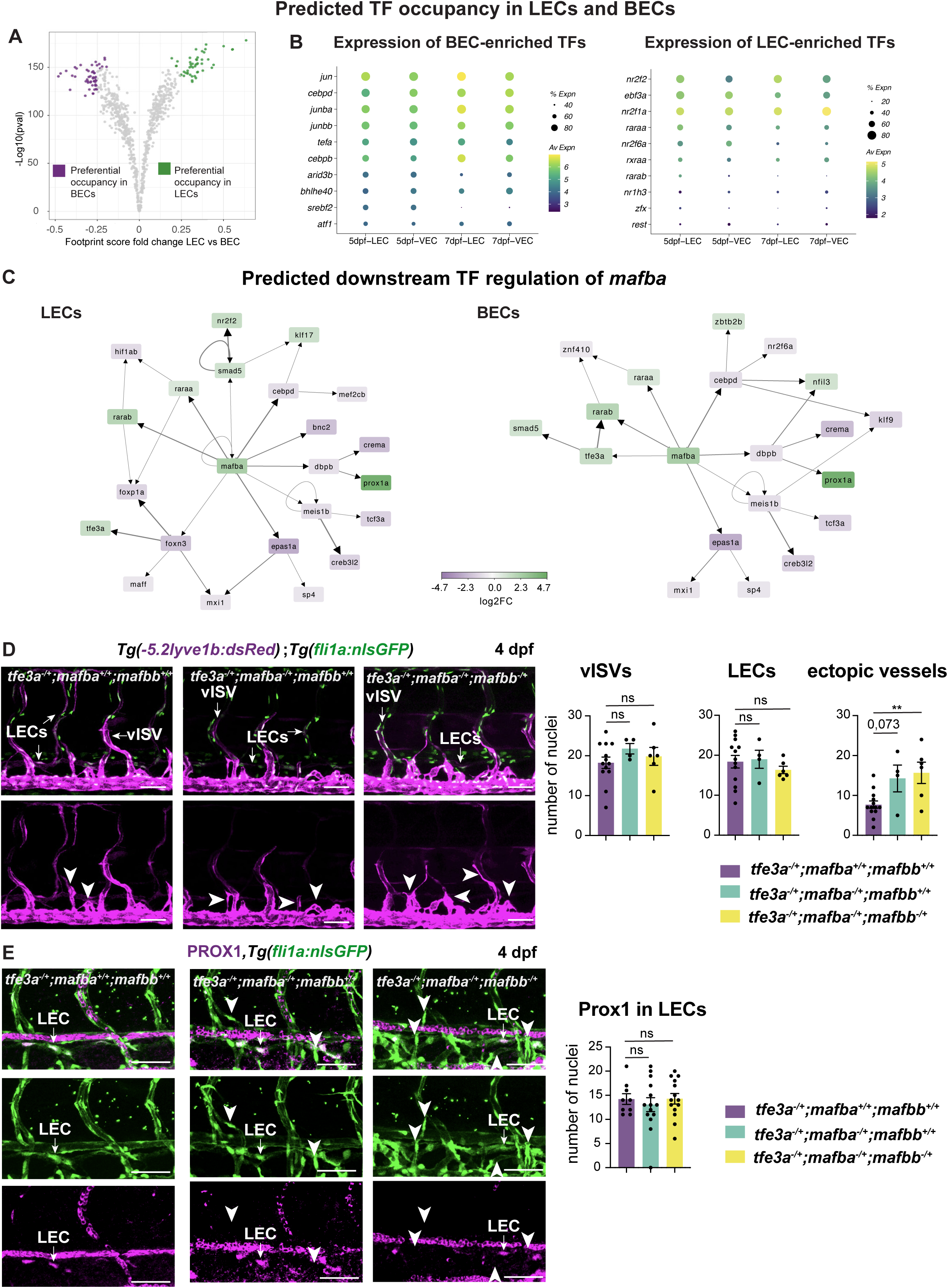
Characterisation of the *mafba* downstream regulatory network in LECs and BECs. (A) Comparison of TF footprints in LECs vs. BECs. Top/bottom 5 percent by score difference is coloured. (B) Dot plots indicating expression of BEC- (left) and LEC-enriched (right) TFs at 5 dpf and 7 dpf. Dot size illustrates percentage of cells presenting transcript sequence counts (% Expn) and viridis scale colour illustrates the average normalised expression (Av Expn) within a cluster. (C) Two-layered TF networks downstream of *mafba* in LECs and BECs, based on the TOBIAS prediction. Green: gene expression enriched in LECs at 5 dpf. Purple: gene expression enriched in BECs at 5 dpf. (D) Quantifications of ectopic vessel formation in triple heterozygous *tfe3a−/+;mafba−/+;mafbb−/+* embryos. Left: Confocal images of *Tg(−5.2lyve1b:dsRed);Tg(fli1a:nlsGFP)* trunk vasculature at 4 dpf in *tfe3a−/+* embryos, *tfe3a−/+;mafba−/+* embryos and *tfe3a−/+;mafba−/+;mafbb−/+* embryos. Arrows: vessel identity. LECs: lymphatic endothelial cells, vISV: venous intersegmental vessels. Arrowheads: ectopic vessels. Right: Quantification of nuclei number in vISV, LECs and ectopic vessels.: ns=no significance, **= p-value 0,0097 (E) Quantification of Prox1-expressing LECs at 4 dpf. Left: immunostaining against Prox1 in the trunk of *Tg(fli1a:nlsGFP) tfe3a−/+* embryos, *tfe3a−/+;mafba−/+* embryos and *tfe3a−/+;mafba−/+;mafbb−/+* embryos. Arrows: Lymphatic vessels. Arrowheads: LECs nuclei. Right: Quantification of PROX1- and GFP-positive nuclei number in LECs. ns=no significance. Scale bars: 50 µm.

To further harness the potential of our dataset, we used the TF footprinting results to investigate the regulatory networks downstream of Mafba, aiming to determine the molecular mechanism by which it regulates lymphatic vessel formation. We identified the promoters bound by Mafba and compared them between LECs and BECs. We identified 1364 predicted sites of occupancy of the Mafba motif at gene promoters in both LECs and BECs, while in 417 sites the motif is predicted to be occupied only in BECs and in 192 sites only in LECs (**Figure S8A, Table S17**). Out of the common targets, 17 genes showed enriched expression in LECs, while 11 were enriched in the VECs at 5 dpf (scRNA-sequencing data with cutoff average log2FC of 0.3, **Figure S8B, Table S18**). Within the promoters predicted to have sites exclusively occupied in LECs, 12 genes showed enriched expression in LECs, and 7 genes showed enriched expression in the VECs at 5 dpf. On the other hand, in the promoters predicted to have sites exclusively occupied in BECs, 0 genes showed enriched expression in LECs and 3 showed enriched expression in VECs at 5 dpf (**Figure S8B, Table S18**). This suggests a complex setup of gene transcription regulatory network, with the likely cooperation of multiple TFs to drive LEC and VEC gene expression and hinting at the potential of Mafba to act as both an activator and repressor.

To fully understand the complexity of Mafba downstream regulation, we built a TF-based two-layered transcriptional network for LECs and BECs (**Figure 7C**). To focus on the differences between LECs and BECs, we included only TFs which were differentially expressed at 5 dpf. In addition to the consensus Mafb motifs from Jaspar, we also considered an endothelial-specific binding motif for *mafba* previously identified in a BEC ChIP-sequencing dataset (Jeong et al., 2017). The number of occupied motif instances in LECs and BECs was lower for the endothelial Mafba motif compared to the Jaspar consensus motifs, with few genes showing enriched expression in LECs and VECs (**Figure S8C-D, Table S19-20**). As a result, the network built on the endothelial motif is composed of only a few genes (**Figure S8E**). This limited predictive ability suggests that this motif might be restricted to BECs. We therefore considered the predictions based on the common motifs going forward, as we were interested in a wider variety of endothelial cells. Noticeably, in both networks we found genes enriched in both LECs or VECs, further supporting the idea that chromatin accessibility does not necessarily indicate active expression (**Figure 7C**). These networks contain known genes involved in vascular and lymphatic development, including members of the retinoic acid signalling family (Bowles et al., 2014), the BMP signalling pathway (Dunworth et al., 2014), and the TFs Prox1 (Koltowska et al., 2015a; Wigle et al., 1999) and Nr2f2 (Srinivasan et al., 2010) (**Figure 7C**).

We have identified the transcriptional signatures characteristic of LECs and BECs, and mapped the downstream regulatory networks of Mafba, uncovering a complex, context-dependent regulatory landscape. The disconnect between lineage-specific gene expression and predicted TF activity highlights the necessity of integrating analysis on multiple layers of gene regulation. Altogether, our multi-omic approach reveals the intricacies of transcriptional control during LEC differentiation and emphasises the critical importance of *in vivo* validation to decode cell-type-specific regulatory programmes.

### Mafba and Tfe3a cooperate to limit ectopic vessel formation

To validate the biological relevance of the identified Mafba transcriptional networks we focused on the *tfe3* gene (**Figure 7C**). This gene has not been previously associated with lymphatic development in zebrafish, but has been shown in mice shown to repress lymphatic fate (Tai-Nagara et al., 2020). Tfe3a also co-operates with Mafb to co-ordinate macrophage differentiation (Zanocco-Marani et al., 2009). *tfe3a* was present in both the LEC and BEC Mafb transcriptional networks: thus, we wanted to investigate its role in the regulation of lymphatic vessel formation. To test if *tfe3a* and *mafba* genetically interact during lymphatic vessel development, we have generated *tfe3a* mutants by introducing 13 bp insertion in exon 5, resulting in a premature stop upstream of the Helix-loop-helix DNA-binding domain 1 (**Figure S8F**). The *tfe3a* mutants grow to adulthood and show no apparent phenotypes. As *mafba* and *mafbb* compensate for each other, and the double-mutants do not develop lymphatic vessels (Arnold et al., 2022) we decided to test the interaction in triple-heterozygous embryos (**Figure 7D**). When comparing *tfe3a* heterozygotes with *tfe3a, mafba* double-heterozygotes or *tfe3a, mafba, mafbb* triple-heterozygotes we noticed no apparent defects in lymphatic or venous vessel formation (**Figure 7D**). Lymphatic vessels differentiate correctly, and show Prox1 expression as well as regular morphology (**Figure 7E)**. Similarly, the venous vessels appear correctly formed and functional (**Figure 7D).** However, in *tfe3a, mafba, mafbb* triple-heterozygous embryos we found a significant increase in ectopic vessels sprouting out of the PCV. Such vessels do not connect to the intersegmental arteries and remain un-luminised (**Figure 7D**). The ectopic vessels did not acquire neither lymphatic identity nor venous functionality. These findings demonstrate that *tfe3a* genetically interacts with *mafba* and *mafbb* to limit ectopic vessel formation and ensure correct vascular differentiation, validating the predictive power of our regulatory networks in uncovering functional genetic interactions shaping vessel formation.

## Discussion

Since the debut of HiC as a powerful tool to investigate chromatin 3D conformation, the consensus has been that TADs are stable structures across different cell types, while the loops they contain shift in a tissue-specific manner to modulate transcriptional activity (Pongubala and Murre, 2021). However, this consensus has recently come under scrutiny, as examples of TADs changing during development (Bonev et al., 2017; Boya et al., 2017; Marti-Marimon et al., 2021; Narendra et al., 2016) or between similar cell lineages (Cresswell and Dozmorov, 2020) emerged. In this work, we have identified a large number of TADs showing differences between our samples, despite LECs and BECs being overall very closely related cell types. These differences indicate that TADs are likely a much more complex feature than initially believed, featuring a nested structure and dynamic changes in their boundaries, and likely play a large role in regulating developmental processes.

Differences in chromatin organisation were also identified at the level of chromatin loops, and the accessible regions they connect. Interestingly, we found that not only the location of loops varies between LECs and BECs, but also that the genomic regions interacting through these loops differ between these two populations. Although the standard model for *cis*-regulation sees the contact of a promoter with an enhancer, different interactions, such as promoter-promoter or enhancer-enhancer have been reported (Li et al., 2012). Promoter-promoter interactions, which in our data are enriched in BECs, have been associated with negative gene regulation, in which one promoter works as a silencer for the other (Wei et al., 2022), while interactions among enhancer elements, enriched in LECs, have a role in driving and coordinating the expression of lineage-specific genes (Madsen et al., 2020). On the other hand, promoter-distal enhancer interactions are more prominent in BECs. Work in *Drosophila* has identified that such interactions are more prominent in differentiated cells (Pollex et al., 2024). As BECs differentiate prior to LECs, we speculate that these differential interactions are an indicator of the differentiated status of these cell types. Overall, these data provide a first look into a more complex level of the regulatory mechanisms active on a genomic level in separating the BEC and LEC lineages.

LECs arise from multiple developmental origins, with progenitors specified from VECs (Küchler et al., 2006; Sabin, 1902; Srinivasan et al., 2007; Wigle et al., 1999; Yaniv et al., 2006) mesenchymal lineage including paraxial mesoderm-derived angioblast (Huntington and McClure, 1910; Lupu et al., 2025), hemogenic endothelium (Stanczuk et al., 2015) or other endothelial sources, such as endothelial angioblast (Eng et al., 2019; Martinez-Corral et al., 2015; Nicenboim et al., 2015; Pichol-Thievend et al., 2018). To differentiate into mature LECs and establish a stable lymphatic identity, these progenitors must coordinate and unify their gene expression programs. This process requires lineage-specific chromatin organisation and precise enhancer-promoter interactions. Previous work in mice and zebrafish has begun to map chromatin accessibility during LECs progenitor specification, mostly focusing on *Prox1* gene regulation and its downstream effectors, and has identified transcription factor signatures marking the lymphatic progenitor pools (Grimm et al., 2023; Kazenwadel et al., 2023; Lupu et al., 2025; Panara et al., 2024). However, to gain a holistic understanding of lymphatic differentiation, gene regulation must be studied at multiple levels. Our goal was to characterise the complexity of gene regulation in LECs, and to define how it differs from that in BECs, a relatively similar cell population. By integrating HiC, bulk ATAC-sequencing and scRNA-sequencing, we uncovered a highly complex regulatory system. Both short- and long-range enhancers are key in driving lymphatic-specific expression of several genes. Moreover, dynamic changes in TADs boundaries selectively isolate or join regulatory elements in a regulatory hub with gene promoters, thereby contributing to the establishment of the LEC molecular identity. These findings lay the foundation for the future studies investigating how chromatin architecture is reorganised during the specification of LEC progenitors, regardless of their cellular origin.

In conclusion, our study provides the first description of overall chromatin organisation in zebrafish LECs and BECs, and validates its connection to differential gene expression *in vivo*. Our datasets provide a powerful tool to further investigate both the developmental mechanism of LEC identity acquisition, and the association of *cis-* dysregulation in lymphatic disease.

## Acknowledgments

This work was supported by Knut och Alice Wallenbergs Stiftelse (2017.0144 and 2022.0187), Ragnar Soderbergs stiftelse (M13/17), Vetenskapsrådet (VR-MH-2016-01437), Kjell och Märta Beijers Stiftelse and Cancerfonden (20 1086 Pj),

The authors acknowledge support from the National Genomics Infrastructure in Stockholm funded by Science for Life Laboratory, the Knut and Alice Wallenberg Foundation and the Swedish Research Council, and SNIC/Uppsala Multidisciplinary Center for Advanced Computational Science for assistance with massively parallel sequencing and access to the UPPMAX computational infrastructure. The authors thank the Bakkers lab for kindly providing the ZED vector.

## Methods

### Zebrafish

Zebrafish work was carried out under ethical approval from the Swedish Board of Agriculture (5.8.18-10590/2018 and 5.8.18-06282/2023). Zebrafish were housed at the Centre for In Vivo Uppsala University (CFVUU, Uppsala, Sweden) and adults and embryos were housed according to the standard laboratory procedure. Published zebrafish lines used were *Tg(fli1a:nEGFP)^y7^*(Lawson and Weinstein, 2002), *TgBAC(prox1a:KalTA4-4xUAS-ADV.E1b:TagRFP)^nim5^*(van Impel et al., 2014), *Tg(5.2lyve1b:Venus)^uu1kk^* (Arnold et al., 2022), *Tg(−5*.*2lyve1b*:*DsRed2)^nz101^*(Okuda et al., 2012)*, Tg(fli1a:H2B-mCherry)^uq37bh^* (Baek et al., 2019), *TgBAC*(*ve-cad:ve-cadTS*)*^uq11bh^* (Lagendijk et al., 2017), *mafba^uq4bh^* (Koltowska et al., 2015b) and *mafbb^uq47bh^* (Arnold et al., 2022). *tfe3a^uu16kk^*, *Tg(+24nrf2f2:basEGFP;ACry:GFP)^uu18kk^, Tg(+47mafba:basEGFP;ACry:GFP)^uu19kk^, Tg(+127tbx1:basEGFP;ACry:GFP)^uu20kk^, Tg(+461mafba:basEGFP;ACry:GFP)^uu21kk^, Tg(−64nrf2f2:basEGFP;ACry:GFP)^uu22kk^, Tg(+41lgmn:basEGFP;ACry:GFP)^uu23kk^* and *Tg(−70mafba:basEGFP;ACry:GFP)^uu24kk^*was generated in this study.

### Cell collection for HiC and bulk ATAC-sequencing

Embryos of *Tg(prox1a:RFP)^nim5^*;*Tg(fli1a:EGFP)^y7^*were collected at 5 days post-fertilisation (dpf). Isolation of cells was performed as previously described (Kartopawiro et al., 2014). For HiC, the crosslinking step of the Arima protocol was performed before sorting. Briefly, the dissociated cells were resuspended in 3% BSA in PBS, then incubated in 1.34% formaldehyde in PBS for 10 minutes at RT. Stop solution 1 was then added as per protocol and the cells incubated for 5 minutes at RT, then 15 minutes on ice. Cells were then washed by repeated centrifugation and resuspension in PBS and used for FACS sorting as described for ATAC-sequencing. The dissociated cells were sorted using a FACS Aria III (BD Biosciences) directly into 300 μl PBS. We based the selection for the desired populations on FSC and SCC, singlets, and alive cells based on their Sytox^TM^ (Thermo Fisher) profile. Double-positive cells were then sorted for the *Tg(prox1a:RFP)^nim5^*and *Tg(fli1a:EGFP)^y7^* transgenes (LECs), whereas single-positive cells were sorted for the *Tg(fli1a:EGFP)^y7^* transgene (BECs), as depicted in **Figure 1A**. At least 50000 cells per population were collected for each repeat. For bulk ATAC-sequencing, cells were then centrifuged and the supernatant removed, after which the tagmentation step was performed as described by the Nextera DNA CD protocol and the material was then snap-froze in dry ice.

### HiC sequencing

Library preparation was performed using an Arima-Hi-C kit (Arima Genomics). Samples were sequenced on NovaSeq6000 (NovaSeq Control Software 1.7.5/RTA v3.4.4) with a 151nt(Read1)-10nt(Index1)-10nt(Index2)-151nt(Read2) setup using ‘NovaSeqXp’ workflow in ‘S4’ mode flowcell.

### Hi-C data analysis and TADs characterisation

Data were processed using Juicer 1.6 (Durand et al., 2016). Reference genome was GRCz11 with alt contigs removed. The restriction map was generated using the script generate_site_positions.py from Juicer version 1.6. The QC metrics and heatmaps of Hi-C contacts are derived from data containing reads aligned with MAPQ ≥ 30. TADs were detected using OnTAD v 1.4 (An et al., 2019), with parameters: “-lsize 7 -penalty 0.075 -maxsz 300” on merged replicate matrices at bin size 10kb. Consensus TAD boundaries were identified using TADCompare (Cresswell and Dozmorov, 2020) interrogating TAD boundaries comparing signal in replicates’ matrices (z_thres=2.5, 500kb window). Non-differential boundaries were considered consensus. TADs delimited by both consensus boundaries were labeled as consensus. Differential boundaries between LEC and BEC were detected using TADCompare (z_thres=2, 500kb window). TADs delimited by at least one differential boundary were labeled differential. For plotting overlapping TADs in Venn and Euler diagrams, TADs with boundaries within 30kb in LECs and BECs were considered equivalent.

### Chromatin loops analysis

Chromatin loops were identified by GPU HiCCUPS (juicer_tools version 2.20.00) using default parameters. Loops from merged_loops.bedpe files were used in further analyses. Loops shared between replicates (tissue consensus loops) within a distance of 25 kb were identified using compare (juicer_tools version 1.9.9). Common and tissue-specific loops were identified by intersecting tissue consensus loops using bedtools v. 2.29.0 (Quinlan and Hall, 2010) (bedtools pairtopair -f 0.4; bedtools pairtopair -type notboth -slop 40000, respectively). We required that common loops overlap at 0.4 of their lengths, and tissue-specific loops are separated by an interval of at least 40 kb.

### Bulk ATAC-sequencing

Library preparation was performed using the Nextera DNA CD kit (Illumina). Samples were sequenced on NextSeq2000 (NextSeq 1000/2000 Control Software 1.2.0.36376/RTA 3.7.17) with a 76nt(Read1)-8nt(Index1)-8nt(Index2)-76nt(Read2) setup using ‘P2’ flowcell.

### Bulk ATAC-sequencing data analysis

Obtained data was processed with the nf-core ATAC-sequencing pipeline v. 1.2.1 (Ewels et al., 2020). Peaks were called using MACS2 in broad mode with FDR cutoff 0.1 (Zhang et al., 2008). Quality control was performed using ATACseqQC (Ou et al., 2018) and peak annotation with Ensembl GRCz11 gene models was performed using ChIPseeker (Yu et al., 2015). Peaks reproducible in at least two out of three replicates were merged to generate a consensus peak set used for differential accessibility analysis, after filtering out features with low counts. Counts were normalised by trimmed mean of M values (TMM) (Robinson and Oshlack, 2010) and differential accessibility (DA) analysis performed using edgeR (Robinson et al., 2010). FDR cutoff of 0.05 was applied to select significant differential peaks.

### Transcription factor footprinting and network construction

TF motifs were downloaded from JASPAR CORE 2022 vertebrate non-redundant collection. For Mafb specific footprints, we also included Mafb motifs from (Jeong et al., 2017) and the Hocomoco MAFB.H12CORE.1.P.B motif (Kulakovskiy et al., 2018). TF site occupancy in consensus peaks was interrogated using TOBIAS (0.16.0) (Bentsen et al., 2020) in differential mode.

TF IDs were mapped to their zebrafish orthologs based on file ORTHOLOGY- ALLIANCE_COMBINED.tsv downloaded from www.alliancegenome.org.

The nodes of the TF networks were constructed from TOBIAS output files based on the annotation of the TF binding site in target promoter (up to 3kb from TSS). The nodes were selected where the target was expressed in 30% of the cells in tissue specific cluster in scRNA-sequencing data, with log2FC > 0.5. The networks were visualised using Cytoscape v. 3.10.1 (Shannon et al., 2003).

### Genomic Visualisations

Composite scaled ATAC-sequencing tracks were generated using deepTools (3.3.2) (Ramírez et al., 2016). The tracks were normalised to 1x average coverage and summarised to produce the average for three replicates. HiC signal was visualised as ICE-corrected merged replicate matrices. Loop coordinates were converted from bedpe format to bed including the outermost bin coordinates. Topmost consensus TADs were visualised on HiC heatmaps as triangles, while three top TAD levels were visualised as intervals, coloured by their TAD score as output by OnTAD. Genomic visualisations were generated using pyGenomeTracks (3.6) (Lopez-Delisle et al., 2021).

### Zebrafish cell dissociation and Fluorescence Activated Cell Sorting (FACS) for scRNA-sequencing

*Tg(−5.2lyve1b:Venus)^uu1kk^;Tg(fli1a:H2B-mCherry)^uq37bh^* embryos were harvested at 5 dpf and 7 dpf. Heads were dissected by making an incision posterior of the otolithic lymph vessel. Trunks were dissected by making an incision posterior of the yolk and an incision posterior of the yolk extension. Isolation of cells was performed by dissociation of the whole embryos, heads and trunks following an optimized published protocol (Kartopawiro et al., 2014). Briefly, we deyolked collected or dissected embryos by pipetting up and down and rinsing in calcium-free ringer’s solution. Then, we centrifuged at 2000 rpm for 5 minutes at 4 °C and removed the supernatant. Then, we dissociated the cells by incubating whole embryos, heads and trunks in 2.5 mg/ml liberase (Sigma-Aldrich) diluted at a 1:35 ratio in DPBS at 28.5 °C. Whole 5 dpf and 7 dpf embryos were incubated 9-10 min, whereas heads and trunks of 5 dpf and 7 dpf embryos were incubated for 7-8 min, homogenizing the samples during and after the incubation. To stop the reaction, we added CaCl2 to a final concentration of 1-2 mM and FBS to a final concentration of 5-10%. We centrifuged at 2000 rpm for 5 minutes at 4 °C, discarding the supernatant. In order to be able to asses live vs. dead cells, we resuspend the cell solution in 1 μM Sytox^TM^ (Thermo Fisher Scientific) in DPBS/EDTA plus 0.5 % FBS. The cell suspension was filtered through a strainer. The dissociated cells were sorted using a FACS Aria III (BD Biosciences) directly into buffered 384- well plates prepared by the Single Cell Core Facility of Flemingsberg Campus (SICOF) on a cold block. We based the selection for the desired populations on FSC and SCC, singlets, and alive cells based on their Sytox^TM^ (Thermo Fisher Scientific) profile. Double-positive cells were sorted for both *Tg(−5.2lyve1b:Venus)^uu1kk^* and *Tg(fli1a:H2B- mCherry)^uq37bh^* transgenes (LECs and VECs), as depicted in **Figure 2A**. After sorting, the 384-well plates were snap-frozen on dry ice and stored at −80 °C before being shipped to the Single Cell Core Facility of Flemingsberg Campus (SICOF).

### scRNA-sequencing

Library preparation and sequencing was performed at the Single Cell Core Facility of Flemingsberg Campus (SICOF). Single-cell cDNA libraries were prepared according to the previously described Smart Seq2 protocol (Vanlandewijck and Betsholtz, 2018). Briefly, mRNA was transcribed into cDNA using oligo(dT) primer and SuperScript II reverse transcriptase (Thermo Fisher Scientific). Second strand cDNA was synthetized using a template switching oligo. The synthetised cDNA was then amplified by polymerase chain reaction (PCR) for 23–26 cycles, depending on the tissue-origin of the respective mRNA sample. Purified cDNA was quality controlled (QC) by analysing on a TapeStation 4200 or 2100 Bioanalyzer with a DNA High Sensitivity chip (Agilent Biotechnologies). When the sample passed the QC, the cDNA was fragmented and tagged (tagmented) using Tn5 transposase, and each single well was uniquely indexed using the Illumina Nextera XT index kits (Set A–D). Thereafter, the uniquely indexed cDNA libraries from one 384-well plate were pooled into one sample to be sequenced on one lane of a HiSeq3000 sequencer (Illumina), using dual indexing and single 50 base-pair reads.

The reads were aligned to the zebrafish genome (GRCz11) and spike-in sequences (from the External RNA Controls Consortium, ERCC92) using STAR v2.5.3a (Dobin et al., 2013), and filtered for uniquely mapping reads. Counts per gene were calculated for each transcript in Ensembl release 97 using rpkmforgenes (Ramsköld et al., 2009). Various QC metrics were generated with RSeQC 2.6.1 (Wang et al., 2012).

### scRNA-sequencing data analysis

Bioinformatics analyses were performed with the R programming language, using Seurat version 5.1.0 (Hao et al., 2021). Reads were normalized to transcripts per million reads (TPMs) in order to compensate for gene length in transcript count estimation. Values were then multiplied by the read length and rounded to the nearest integer. Cell libraries were considered of low quality and filtered out if: (1) the percentages of mitochondrial and ribosomal gene families were above 25 %; (2) the percentage of protein-coding genes was below 90 %; (3) the number of unique genes in a cell was below two standard deviations from the mean of the plate (**Figure S2A- B)**. After filtering, the datasets contained 1264 cells for 5 dpf, and 1139 cells for 7 dpf. Genes detected in less than three cells were excluded from the analysis. For further downstream analysis only protein coding genes were used.

To remove batch differences, the Seurat SCT integration (Hafemeister C and Satija R, 2019; Stuart T et al., 2019) pipeline was used. The dataset was divided based on the different anatomical parts of the zebrafish. Each subset was normalized individually and variable features were identified. Based on these features, integration anchors were derived and used for integrating the subsets into one dataset used for subsequent analysis. Similarly, integration was performed for sub-divisions of the dataset for the different developmental time points individually.

The datasets were further processed for identification of highly variable features and linear data compression using principal component analysis (PCA). Non-linear dimensionality reduction was run on the top 50 PCs to obtain an UMAP embedded. Unsupervised graph clustering was run using the Louvain algorithm with the modularity resolution parameter (res = 0.5). Cluster marker genes were identified using the Wilcoxon rank-sum test. Genes with average logarithmic fold change of expression (log_2_FC) > 0.25 at Bonferroni adjusted p-value < 0.01 were considered.

For all scRNA-sequencing datasets, exclusion of contamination clusters, as well as identification of the relevant LEC and VEC clusters was determined using DEG and the expression patterns of the described markers. All downstream analysis and visualization were performed using Seurat version 5.1.0 (Hao et al., 2021) with default settings, and utilizing the ggplot2 (“ggplot2: Elegant Graphics for Data Analysis | SpringerLink,” 2024) R package and enrichplot (“Yu G. enrichplot: Visualization of Functional Enrichment Result. R package version 1.24.2,” 2024) Bioconductor package.

### Enhancer identification

Short-range candidate enhancers were identified as peaks of open chromatin in LECs in the vicinity of genes whose expression is enriched in LECs at either 5 dpf or 7 dpf. Long-range candidate enhancers were defined as differentially accessible regions of non-coding DNA which interacts with the open promoter of a LEC-enriched gene through a LEC-specific loop. First, Rstudio 2023.12.1+402 (Posit team, 2024; R Core Team, 2023) was used to extract the coordinates of the left and right ends of the LEC-specific loops. Coordinates of the accessible TSS in LECs and the LEC-specific intronic and distal accessible regions were obtained in the same way. Using bedops v2.4.41 (Neph et al., 2012) with the –closest function, the closest open TSS and LEC-specific distal or intronic open elements to the ends of the loops were identified. In order to ensure their relation to the loop, only those located at less than 10kb from the loop ends were further considered in the analysis. Using Rstudio, loops marked by an accessible TSS on one end and a LEC-specific accessible distal or intronic element on the other were identified. Then, the data was further refined by intersecting this list with that of DE genes in LECs at 5 dpf and 7 dpf.

### Enhancer conservation analysis

Sequence conservation was tested using mVISTA (Brudno et al., 2003; Dubchak et al., 2000; Frazer et al., 2004; Mayor et al., 2000) as previously described (Panara et al., 2024). The sequences were oriented in the direction of the transcription. The species, assemblies and regions used are listed in **Supplementary Table 21**.

### Generation of transgenic lines

To test the activity of short-range enhancers, the elements of interest were ether cloned into the ZED vector as previously described (Bessa et al., 2009). For long-range enhancers, reporter vectors using a minimal E1b promoter as previously described (Panara et al., 2024) were ordered from GenScript. Sequences are in the **Supplementary Table 22**.

To generate mosaic transgenic lines, 1μl of construct at 20 ng/μl and tol2 transposase mRNA at 100 ng/μl was injected into the one-cell stage *Tg(prox1a:RFP)^nim5^* zebrafish embryos. Embryos were imaged in F0. At least 4 embryos per round of injection were imaged (**Supplementary Table 23**).

To generate in F1 transgenic lines and assess the enhancer activity, 1μl of construct at 20 ng/μl and tol2 transposase mRNA at 100 ng/μl was injected into the one-cell stage *Tg(prox1a:RFP)^nim5^* zebrafish embryos and raised to adulthoods. The F0 adults were outcrossed to *Tg(prox1a:RFP)^nim5^*and F1 embryos were screened for expression. At least 2 injected adult F0 were screened per enhancer. Embryos with positive green lenses were imaged using confocal microscope to determine enhancer-driven expression (**Supplementary Table 23**).

### Tfe3a CRISPR mutant generation

*Tfe3a* mutants were generated using the IDT Alt-R^TM^ CRISPR-Cas9 System. The guide was designed using IDT’s Predesigned Alt-R^TM^ CRISPR-Cas9 guide RNA Design Tool in the exon 5 of the *tfe3a* gene. Primers were designed upstream and downstream of the guide site as described in the protocol originally developed in Burgess Lab (Carrington et al., 2015) and adapted for the DanioReadout facility (Habicher et al., 2022).

Zebrafish embryos were injected at the one-cell stage with 50-100 ng/μL of gRNA complex (crRNA plus tracrRNA) and 200ng/μL Alt-R Cas9 Nuclease V3. F0 fish were raised to adulthood and outcrossed to wildtype. F1 embryos from three individual crosses were raised. F1 fin clips were genotyped using Fragment Length Analysis. A founder containing an insertion of 13 bp was confirmed by Sanger Sequencing and a stable line was established.

Guide Sequence: ATTTATGACGCTGATCGACT

Genotyping primers used (with M13 tag and PIGtail tag in capital letters)

Forward Primer Sequence: TGTAAAACGACGGCCAGccagctaagccgttcgttct

Reverse Primer Sequence: GTGTCTTgggtgaactatcccttccaagca

### In situ probe synthesis

Probes were generated by In-Fusion® HD cloning (Takara Bio) to the PCS2+ vector using the primers listed in **Supplementary Table 24**. Vectors were then linearised with BalmHI for antisense probes and SnaBI for sense probes prior to probe synthesis was then carried out using the MAXIscript T7 Transcrption Kit (Invitrogen).

### In situ hybridisation

Embryos of the desired stage were dechorionated prior to fixation in 4% paraformaldehyde/PBS for 3 hours at room temperature or overnight at 4°C. Fixed embryos were dehydrated with progressive washed of 50% MeOH, 75% MeOH and 100% MeOH then stored in 100% methanol at −20°C. Embryos at 2 dpf were incubated in 10mg/ml PK in PBST for 30 minutes at RT and embryos at 5 dpf were incubated in 20mg/ml PK in PBST for 30 minutes at RT. Subsequently, embryos were fixed in 4% PFA for 20 minutes prior to processing by *in situ* hybridisation as previously described by (Xu et al., 1994) with BCIP/NBT. After colour development embryos were fixed in 4% PFA for 20 minutes then cleared in 75% glycerol/PBS, and mounted for imaging using a Leica Fluorescent Stereo Microscope M165 FC and 1x objective (objective number: 10450028). Embryos were mounted in 3% methylcellulose.

### Fluorescent in situ hybridisation

The protocol for standard *in situ* hybridisation was followed until the blocking step. From here, embryos were blocked in 2 mg/ml BSA (Sigma)/2% sheep serum (Thermo Fisher) for 3 hours, prior to antibody incubation in 1:5000 anti-digoxigenin fluorescein POD (Roche) and 1:400 Chicken Anti-GFP (Abcam) at 4°C for 16 hours. Embryos were washed 8 times in PBST for 30 minutes prior to incubation with 1:100 tyramide- Cy3 in 1x amplification diluent (Perkin Elmer). Embryos were washed a further 8 times in PBST for 30 minutes prior to blocking in 2 mg/ml BSA (Sigma)/2% sheep serum (Thermo Fisher) for a minimum of 1 hour. Embryos were incubated in 1:400 Anti-chicken 488 (Jackson Immuno Research) at 4°C for 16 hours. Embryos were then washed 8 times in PBST for 15 minutes prior to fixation in 4% PFA for 30 minutes. Embryos were imaged on a Leica TCS SP8 DLS with HC PL APO CS2 40× water objective (objective number: 11506360).

### Immunostainings

Embryos were fixed at 4 dpf overnight at 4°C and a previously described protocol was followed (Koltowska et al., 2015a; Shin et al., 2002). We applied the following modification: the Proteinase K treatment was performed at 20μg/ml for 35 minutes and blocking prior to primary antibody was done for 48 hours. For primary antibodies Chicken anti-GFP (1:400, Abcam) and Rabbit anti-Prox1 (1:500 AngioBio Co.) were used and for secondary antibodies Anti-Rabbit IgG-HRP (1:1000 CellSignalling Technology) and Anti-Chicken Alexa-488 (1:200 703-545-155 Jackson Immuno Research)) followed by the TSA™ Plus Cyanine 3 System amplification according to the manufacturer instructions (NEL744001KT Perkin Elmer) and DAPI (1:1000 D1306 Thermo Fisher Scientific) staining. The trunks were mounted in the clearing solution Omnipaque (350 mg litre^−1^ concentration per 1 ml iohexol, GE Healthcare) for imaging.

### Imaging, image processing and quantification

Embryos for *in vivo* fluorescent imaging were anesthetised with tricaine and mounted in 1% low melting agarose. Embryos were imaged using a Leica TCS SP8 DLS inverted confocal microscope with a Fluotar VISR 25X water objective (objective number: 11506375). The 488 nm (for GFP) and 552 nm (for RFP or DsRed) laser lines were used. For trunk vasculature the images were taken between 8-15 somites across the width of the fish, generating a full z-stack. For facial lymphatic vessels, whole z-stack consisted of the vessels on one side of the fish.

Masking was performed in Imaris v9.3.0 with a surface detail of 1μm. Images were processed using ImageJ 2.9.0. All representative images are Maximum Intensity Projections.

Manual quantification of lymphatic vessels, vIVS and ectopic vessel cell numbers was performed using full z-stacks. The number was counted using the overlay of DsRed- and GFP-channels over two somites in the trunks. The vessel type was distinguished based on the morphology (Okuda et al., 2012). Quantification was performed by scoring the amount of nuclei from corresponding vessel types for each genotype, as previously described (Arnold et al., 2022). Quantification of Prox1-positive lymphatic endothelial cell was conducted using a full z-stack of trunk vessels, based on the co-presence of *nfli:GFP* and Prox1 signal. The double-positive GFP and Prox1 nuclei were recorded for thoracic duct and intersegmental lymphatic vessels.

### Statistical analysis

Fisher’s test was used to investigate the distribution of differential TADs in the proximity of LEC and BEC enriched loci (**Figure 2D**).

For loop interaction classification (**Figure S5B-D**), the expected distribution of features in loops was calculated based on the frequency of the features in either the open chromatin peaks or the differentially accessible peaks from bulk ATAC-sequencing analysis. The expected distribution was compared with the observed one with a chi-square test for goodness of fit using Prism 9 (Graphpad). Observed distributions in LECs and BECs were compared using a chi-square test. Residuals were calculated by computing the expected number of loops in each category and subtracting the value from the observed count. For multiple group comparisons (**Figure 7D-E**) the normality was tested with a Shapiro-Wilk test, all quantification but ectopic vessels passed the normality test. The ordinary one-way ANOVA test was used for normally distributed data, Kruskal-Wallis test was used for non-normally distributed data. All error bars represent standard error of the mean (SEM).

### Resource availability

Upon publication the Hi-C, scRNA-sequencing and ATAC-sequencing data will be made publicly available.

## Bibliography

An, L., Yang, T., Yang, J., Nuebler, J., Xiang, G., Hardison, R.C., Li, Q., Zhang, Y., 2019. OnTAD: hierarchical domain structure reveals the divergence of activity among TADs and boundaries. Genome Biol 20, 282. 10.1186/s13059-019-1893-y

Arnold, H., Panara, V., Hußmann, M., Filipek-Gorniok, B., Skoczylas, R., Ranefall, P., Gloger, M., Allalou, A., Hogan, B.M., Schulte-Merker, S., Koltowska, K., 2022. mafba and mafbb differentially regulate lymphatic endothelial cell migration in topographically distinct manners. Cell Reports 39, 110982. 10.1016/j.celrep.2022.110982

Baek, S., Oh, T.G., Secker, G., Sutton, D.L., Okuda, K.S., Paterson, S., Bower, N.I., Toubia, J., Koltowska, K., Capon, S.J., Baillie, G.J., Simons, C., Muscat, G.E.O., Lagendijk, A.K., Smith, K.A., Harvey, N.L., Hogan, B.M., 2019. The Alternative Splicing Regulator Nova2 Constrains Vascular Erk Signaling to Limit Specification of the Lymphatic Lineage. Developmental cell 49, 279–292.e5. 10.1016/j.devcel.2019.03.017

Bentsen, M., Goymann, P., Schultheis, H., Klee, K., Petrova, A., Wiegandt, R., Fust, A., Preussner, J., Kuenne, C., Braun, T., Kim, J., Looso, M., 2020. ATAC-seq footprinting unravels kinetics of transcription factor binding during zygotic genome activation. Nat Commun 11, 4267. 10.1038/s41467-020-18035-1

Bessa, J., Tena, J.J., de la Calle-Mustienes, E., Fernández-Miñán, A., Naranjo, S., Fernández, A., Montoliu, L., Akalin, A., Lenhard, B., Casares, F., Gómez-Skarmeta, J.L., 2009. Zebrafish enhancer detection (ZED) vector: A new tool to facilitate transgenesis and the functional analysis of *cis* -regulatory regions in zebrafish. Dev. Dyn. 238, 2409– 2417. 10.1002/dvdy.22051

Bonev, B., Mendelson Cohen, N., Szabo, Q., Fritsch, L., Papadopoulos, G.L., Lubling, Y., Xu, X., Lv, X., Hugnot, J.-P., Tanay, A., Cavalli, G., 2017. Multiscale 3D Genome Rewiring during Mouse Neural Development. Cell 171, 557–572.e24. 10.1016/j.cell.2017.09.043

Bowles, J., Secker, G., Nguyen, C., Kazenwadel, J., Truong, V., Frampton, E., Curtis, C., Skoczylas, R., Davidson, T.-L., Miura, N., Hong, Y.-K., Koopman, P., Harvey, N.L., François, M., 2014. Control of retinoid levels by CYP26B1 is important for lymphatic vascular development in the mouse embryo. Developmental Biology 386, 25–33. 10.1016/j.ydbio.2013.12.008

Boya, R., Yadavalli, A.D., Nikhat, S., Kurukuti, S., Palakodeti, D., Pongubala, J.M.R., 2017. Developmentally regulated higher-order chromatin interactions orchestrate B cell fate commitment. Nucleic Acids Research 45, 11070–11087. 10.1093/nar/gkx722

Brudno, M., Do, C.B., Cooper, G.M., Kim, M.F., Davydov, E., Green, E.D., Sidow, A., Batzoglou, S., 2003. LAGAN and Multi-LAGAN: efficient tools for large-scale multiple alignment of genomic DNA. Genome Res 13, 721–731. 10.1101/gr.926603

Carrington, B., Varshney, G.K., Burgess, S.M., Sood, R., 2015. CRISPR-STAT: an easy and reliable PCR-based method to evaluate target-specific sgRNA activity. Nucleic Acids Res 43, e157. 10.1093/nar/gkv802

Chen, L., Mupo, A., Huynh, T., Cioffi, S., Woods, M., Jin, C., McKeehan, W., Thompson-Snipes, L., Baldini, A., Illingworth, E., 2010. Tbx1 regulates Vegfr3 and is required for lymphatic vessel development. The Journal of Cell Biology 189, 417–424. 10.1083/jcb.200912037

Cresswell, K.G., Dozmorov, M.G., 2020. TADCompare: An R Package for Differential and Temporal Analysis of Topologically Associated Domains. Front. Genet. 11, 158. 10.3389/fgene.2020.00158

Dobin, A., Davis, C.A., Schlesinger, F., Drenkow, J., Zaleski, C., Jha, S., Batut, P., Chaisson, M., Gingeras, T.R., 2013. STAR: ultrafast universal RNA-seq aligner. Bioinformatics 29, 15–21. 10.1093/bioinformatics/bts635

Dubchak, I., Brudno, M., Loots, G.G., Pachter, L., Mayor, C., Rubin, E.M., Frazer, K.A., 2000. Active conservation of noncoding sequences revealed by three-way species comparisons. Genome Res 10, 1304–1306. 10.1101/gr.142200

Dunworth, W.P., Cardona-Costa, J., Bozkulak, E.C., Kim, J.-D., Meadows, S., Fischer, J.C., Wang, Y., Cleaver, O., Qyang, Y., Ober, E.A., Jin, S.-W., 2014. Bone Morphogenetic Protein 2 Signaling Negatively Modulates Lymphatic Development in Vertebrate Embryos. Circulation Research 114, 56–66. 10.1161/CIRCRESAHA.114.302452

Durand, N.C., Shamim, M.S., Machol, I., Rao, S.S.P., Huntley, M.H., Lander, E.S., Aiden, E.L., 2016. Juicer Provides a One-Click System for Analyzing Loop-Resolution Hi-C Experiments. Cell Systems 3, 95–98. 10.1016/j.cels.2016.07.002

Eng, T.C., Chen, W., Okuda, K.S., Misa, J.P., Padberg, Y., Crosier, K.E., Crosier, P.S., Hall, C.J., Schulte-Merker, S., Hogan, B.M., Astin, J.W., 2019. Zebrafish facial lymphatics develop through sequential addition of venous and non-venous progenitors. EMBO Reports 20, e47079. 10.15252/embr.201847079

Ewels, P.A., Peltzer, A., Fillinger, S., Patel, H., Alneberg, J., Wilm, A., Garcia, M.U., Di Tommaso, P., Nahnsen, S., 2020. The nf-core framework for community-curated bioinformatics pipelines. Nat Biotechnol 38, 276–278. 10.1038/s41587-020-0439-x

François, M., Caprini, A., Hosking, B., Orsenigo, F., Wilhelm, D., Browne, C., Paavonen, K., Karnezis, T., Shayan, R., Downes, M., Davidson, T., Tutt, D., Cheah, K.S.E., Stacker, S.A., Muscat, G.E.O., Achen, M.G., Dejana, E., Koopman, P., 2008. Sox18 induces development of the lymphatic vasculature in mice. Nature 456, 643–647. 10.1038/nature07391

Franke, M., De La Calle-Mustienes, E., Neto, A., Almuedo-Castillo, M., Irastorza-Azcarate, I., Acemel, R.D., Tena, J.J., Santos-Pereira, J.M., Gómez-Skarmeta, J.L., 2021. CTCF knockout in zebrafish induces alterations in regulatory landscapes and developmental gene expression. Nat Commun 12, 5415. 10.1038/s41467-021-25604-5

Frazer, K.A., Pachter, L., Poliakov, A., Rubin, E.M., Dubchak, I., 2004. VISTA: computational tools for comparative genomics. Nucleic Acids Res 32, W273–279. 10.1093/nar/gkh458

Gauvrit, S., Villasenor, A., Strilic, B., Kitchen, P., Collins, M.M., Marín-Juez, R., Guenther, S., Maischein, H.-M., Fukuda, N., Canham, M.A., Brickman, J.M., Bogue, C.W., Jayaraman, P.-S., Stainier, D.Y.R., 2018. HHEX is a transcriptional regulator of the VEGFC/FLT4/PROX1 signaling axis during vascular development. Nat Commun 9, 2704. 10.1038/s41467-018-05039-1

ggplot2: Elegant Graphics for Data Analysis | SpringerLink [WWW Document], 2024. URL https://link.springer.com/book/10.1007/978-3-319-24277-4 (accessed 8.22.24).

Grimm, L., Mason, E., Yu, H., Dudczig, S., Panara, V., Chen, T., Bower, N.I., Paterson, S., Rondon Galeano, M., Kobayashi, S., Senabouth, A., Lagendijk, A.K., Powell, J., Smith, K.A., Okuda, K.S., Koltowska, K., Hogan, B.M., 2023. Single-cell analysis of lymphatic endothelial cell fate specification and differentiation during zebrafish development. The EMBO Journal n/a, e112590. 10.15252/embj.2022112590

Habicher, J., Varshney, G.K., Waldmann, L., Snitting, D., Allalou, A., Zhang, H., Ghanem, A., Öhman Mägi, C., Dierker, T., Kjellén, L., Burgess, S.M., Ledin, J., 2022. Chondroitin/dermatan sulfate glycosyltransferase genes are essential for craniofacial development. PLoS Genet 18, e1010067. 10.1371/journal.pgen.1010067

Hafemeister C, Satija R, 2019. Normalization and variance stabilization of single-cell RNA-seq data using regularized negative binomial regression. Genome biology 20. 10.1186/s13059-019-1874-1

Hao, Y., Hao, S., Andersen-Nissen, E., Mauck, W.M., Zheng, S., Butler, A., Lee, M.J., Wilk, A.J., Darby, C., Zager, M., Hoffman, P., Stoeckius, M., Papalexi, E., Mimitou, E.P., Jain, J., Srivastava, A., Stuart, T., Fleming, L.M., Yeung, B., Rogers, A.J., McElrath, J.M., Blish, C.A., Gottardo, R., Smibert, P., Satija, R., 2021. Integrated analysis of multimodal single-cell data. Cell 184, 3573–3587.e29. 10.1016/j.cell.2021.04.048

Hernández Vásquez, M.N., Ulvmar, M.H., González-Loyola, A., Kritikos, I., Sun, Y., He, L., Halin, C., Petrova, T.V., Mäkinen, T., 2021. Transcription factor FOXP2 is a flow-induced regulator of collecting lymphatic vessels. The EMBO Journal 40, e107192. 10.15252/embj.2020107192

Hu, Z., Zhao, X., Wu, Z., Qu, B., Yuan, M., Xing, Y., Song, Y., Wang, Z., 2024. Lymphatic vessel: Origin, heterogeneity, biological functions and therapeutic targets. Signal Transduction and Targeted Therapy 9, 9. 10.1038/s41392-023-01723-x

Huntington, G.S., McClure, C.F.W., 1910. The anatomy and development of the jugular lymph sacs in the domestic cat (Felis domestica). Am. J. Anat. 10, 177–312. 10.1002/aja.1000100108

Jeong, H.-W., Hernández-Rodríguez, B., Kim, J., Kim, K.-P., Enriquez-Gasca, R., Yoon, J., Adams, S., Schöler, H.R., Vaquerizas, J.M., Adams, R.H., 2017. Transcriptional regulation of endothelial cell behavior during sprouting angiogenesis. Nature Communications 1–14. 10.1038/s41467-017-00738-7

Kartopawiro, J., Bower, N.I., Karnezis, T., Kazenwadel, J., Betterman, K.L., Lesieur, E., Koltowska, K., Astin, J., Crosier, P., Vermeren, S., Achen, M.G., Stacker, S.A., Smith, K.A., Harvey, N.L., François, M., Hogan, B.M., 2014. Arap3 is dysregulated in a mouse model of hypotrichosis-lymphedema-telangiectasia and regulates lymphatic vascular development. Hum Mol Genet 23, 1286–1297. 10.1093/hmg/ddt518

Kazenwadel, J., Betterman, K.L., Chong, C.-E., Stokes, P.H., Lee, Y.K., Secker, G.A., Agalarov, Y., Demir, C.S., Lawrence, D.M., Sutton, D.L., Tabruyn, S.P., Miura, N., Salminen, M., Petrova, T.V., Matthews, J.M., Hahn, C.N., Scott, H.S., Harvey, N.L., 2015. GATA2 is required for lymphatic vessel valve development and maintenance. J. Clin. Invest. 125, 2979–2994. 10.1172/JCI78888

Kazenwadel, J., Venugopal, P., Oszmiana, A., Toubia, J., Arriola-Martinez, L., Panara, V., Piltz, S.G., Brown, C., Ma, W., Schreiber, A.W., Koltowska, K., Taoudi, S., Thomas, P.Q., Scott, H.S., Harvey, N.L., 2023. A Prox1 enhancer represses haematopoiesis in the lymphatic vasculature. Nature 614, 343–348. 10.1038/s41586-022-05650-9

Koltowska, K., Lagendijk, A.K., Pichol-Thievend, C., Fischer, J.C., Francois, M., Ober, E.A., Yap, A.S., Hogan, B.M., 2015a. Vegfc Regulates Bipotential Precursor Division and Prox1 Expression to Promote Lymphatic Identity in Zebrafish. Cell Reports 13, 1828– 1841. 10.1016/j.celrep.2015.10.055

Koltowska, K., Paterson, S., Bower, N.I., Baillie, G.J., Lagendijk, A.K., Astin, J.W., Chen, H., François, M., Crosier, P.S., Taft, R.J., Simons, C., Smith, K.A., Hogan, B.M., 2015b. mafbais a downstream transcriptional effector of Vegfc signaling essential for embryonic lymphangiogenesis in zebrafish. Genes & Development 29, 1618–1630. 10.1101/gad.263210.115

Küchler, A.M., Gjini, E., Peterson-Maduro, J., Cancilla, B., Wolburg, H., Schulte-Merker, S., 2006. Development of the Zebrafish Lymphatic System Requires Vegfc Signaling. Current Biology 16, 1244–1248. 10.1016/j.cub.2006.05.026

Kulakovskiy, I.V., Vorontsov, I.E., Yevshin, I.S., Sharipov, R.N., Fedorova, A.D., Rumynskiy, E.I., Medvedeva, Y.A., Magana-Mora, A., Bajic, V.B., Papatsenko, D.A., Kolpakov, F.A., Makeev, V.J., 2018. HOCOMOCO: towards a complete collection of transcription factor binding models for human and mouse via large-scale ChIP-Seq analysis. Nucleic Acids Res 46, D252–D259. 10.1093/nar/gkx1106

Lagendijk, A.K., Gomez, G.A., Baek, S., Hesselson, D., Hughes, W.E., Paterson, S., Conway, D.E., Belting, H.-G., Affolter, M., Smith, K.A., Schwartz, M.A., Yap, A.S., Hogan, B.M., 2017. Live imaging molecular changes in junctional tension upon VE-cadherin in zebrafish. Nature Communications 8, 1–12. 10.1038/s41467-017-01325-6

Lawson, N.D., Weinstein, B.M., 2002. In vivo imaging of embryonic vascular development using transgenic zebrafish. Developmental Biology 248, 307–318.

Li, G., Ruan, X., Auerbach, R.K., Sandhu, K.S., Zheng, M., Wang, P., Poh, H.M., Goh, Y., Lim, J., Zhang, J., Sim, H.S., Peh, S.Q., Mulawadi, F.H., Ong, C.T., Orlov, Y.L., Hong, S., Zhang, Z., Landt, S., Raha, D., Euskirchen, G., Wei, C.-L., Ge, W., Wang, H., Davis, C., Fisher-Aylor, K.I., Mortazavi, A., Gerstein, M., Gingeras, T., Wold, B., Sun, Y., Fullwood, M.J., Cheung, E., Liu, E., Sung, W.-K., Snyder, M., Ruan, Y., 2012. Extensive promoter-centered chromatin interactions provide a topological basis for transcription regulation. Cell 148, 84–98. 10.1016/j.cell.2011.12.014

Lopez-Delisle, L., Rabbani, L., Wolff, J., Bhardwaj, V., Backofen, R., Grüning, B., Ramírez, F., Manke, T., 2021. pyGenomeTracks: reproducible plots for multivariate genomic datasets. Bioinformatics 37, 422–423. 10.1093/bioinformatics/btaa692

Lupu, I.-E., Grainger, D.E., Kirschnick, N., Weischer, S., Zhao, E., Martinez-Corral, I., Schoofs, H., Vanhollebeke, M., Jones, G., Godwin, J., Forrow, A., Lahmann, I., Riley, P.R., Zobel, T., Alitalo, K., Mäkinen, T., Kiefer, F., Stone, O.A., 2025. Direct specification of lymphatic endothelium from mesenchymal progenitors. Nat Cardiovasc Res 4, 45–63. 10.1038/s44161-024-00570-5

Madsen, J.G.S., Madsen, M.S., Rauch, A., Traynor, S., Van Hauwaert, E.L., Haakonsson, A.K., Javierre, B.M., Hyldahl, M., Fraser, P., Mandrup, S., 2020. Highly interconnected enhancer communities control lineage-determining genes in human mesenchymal stem cells. Nat Genet 52, 1227–1238. 10.1038/s41588-020-0709-z

Mañes-García, J., Marco-Ferreres, R., Beccari, L., 2024. Shaping gene expression and its evolution by chromatin architecture and enhancer activity, in: Current Topics in Developmental Biology. Elsevier, pp. 406–437. 10.1016/bs.ctdb.2024.01.001

Marti-Marimon, M., Vialaneix, N., Lahbib-Mansais, Y., Zytnicki, M., Camut, S., Robelin, D., Yerle-Bouissou, M., Foissac, S., 2021. Major Reorganization of Chromosome Conformation During Muscle Development in Pig. Front. Genet. 12, 748239. 10.3389/fgene.2021.748239

Martinez-Corral, I., Ulvmar, M.H., Stanczuk, L., Tatin, F., Kizhatil, K., John, S.W.M., Alitalo, K., Ortega, S., Makinen, T., 2015. Nonvenous Origin of Dermal Lymphatic Vasculature. Circulation Research 116, 1649–1654. 10.1161/CIRCRESAHA.116.306170

Mayor, C., Brudno, M., Schwartz, J.R., Poliakov, A., Rubin, E.M., Frazer, K.A., Pachter, L.S., Dubchak, I., 2000. VISTA : visualizing global DNA sequence alignments of arbitrary length. Bioinformatics 16, 1046–1047. 10.1093/bioinformatics/16.11.1046

Narendra, V., Bulajić, M., Dekker, J., Mazzoni, E.O., Reinberg, D., 2016. CTCF-mediated topological boundaries during development foster appropriate gene regulation. Genes Dev. 30, 2657–2662. 10.1101/gad.288324.116

Neph, S., Kuehn, M.S., Reynolds, A.P., Haugen, E., Thurman, R.E., Johnson, A.K., Rynes, E., Maurano, M.T., Vierstra, J., Thomas, S., Sandstrom, R., Humbert, R., Stamatoyannopoulos, J.A., 2012. BEDOPS: high-performance genomic feature operations. Bioinformatics 28, 1919–1920. 10.1093/bioinformatics/bts277

Nicenboim, J., Malkinson, G., Lupo, T., Asaf, L., Sela, Y., Mayseless, O., Gibbs-Bar, L., Senderovich, N., Hashimshony, T., Shin, M., Jerafi-Vider, A., Avraham-Davidi, I., Krupalnik, V., Hofi, R., Almog, G., Astin, J.W., Golani, O., Ben-Dor, S., Crosier, P.S., Herzog, W., Lawson, N.D., Hanna, J.H., Yanai, I., Yaniv, K., 2015. Lymphatic vessels arise from specialized angioblasts within a venous niche. Nature 522, 56–61. 10.1038/nature14425

Norrmén, C., Ivanov, K.I., Cheng, J., Zangger, N., Delorenzi, M., Jaquet, M., Miura, N., Puolakkainen, P., Horsley, V., Hu, J., Augustin, H.G., Ylä-Herttuala, S., Alitalo, K., Petrova, T.V., 2009. FOXC2 controls formation and maturation of lymphatic collecting vessels through cooperation with NFATc1. Journal of Cell Biology 185, 439–457. 10.1083/jcb.200901104

Okuda, K.S., Astin, J.W., Misa, J.P., Flores, M.V., Crosier, K.E., Crosier, P.S., 2012. lyve1 expression reveals novel lymphatic vessels and new mechanisms for lymphatic vessel development in zebrafish. Development 139, 2381–2391. 10.1242/dev.077701

Ou, J., Liu, H., Yu, J., Kelliher, M.A., Castilla, L.H., Lawson, N.D., Zhu, L.J., 2018. ATACseqQC: a Bioconductor package for post-alignment quality assessment of ATAC-seq data. BMC Genomics 19, 169. 10.1186/s12864-018-4559-3

Panara, V., Yu, H., Peng, D., Staxäng, K., Hodik, M., Filipek-Gorniok, B., Kazenwadel, J., Skoczylas, R., Mason, E., Allalou, A., Harvey, N.L., Haitina, T., Hogan, B.M., Koltowska, K., 2024. Multiple cis-regulatory elements control prox1a expression in distinct lymphatic vascular beds. Development 151, dev202525. 10.1242/dev.202525

Payne, S., Neal, A., De Val, S., 2024. Transcription factors regulating vasculogenesis and angiogenesis. Developmental Dynamics 253, 28–58. 10.1002/dvdy.575

Petrova, T.V., Karpanen, T., Norrmén, C., Mellor, R., Tamakoshi, T., Finegold, D., Ferrell, R., Kerjaschki, D., Mortimer, P., Ylä-Herttuala, S., Miura, N., Alitalo, K., 2004. Defective valves and abnormal mural cell recruitment underlie lymphatic vascular failure in lymphedema distichiasis. Nat Med 10, 974–981. 10.1038/nm1094

Pichol-Thievend, C., Betterman, K.L., Liu, X., Ma, W., Skoczylas, R., Lesieur, E., Bos, F.L., Schulte, D., Schulte-Merker, S., Hogan, B.M., Oliver, G., Harvey, N.L., Francois, M., 2018. A blood capillary plexus-derived population of progenitor cells contributes to genesis of the dermal lymphatic vasculature during embryonic development. Development 145, dev160184. 10.1242/dev.160184

Pollex, T., Rabinowitz, A., Gambetta, M.C., Marco-Ferreres, R., Viales, R.R., Jankowski, A., Schaub, C., Furlong, E.E.M., 2024. Enhancer–promoter interactions become more instructive in the transition from cell-fate specification to tissue differentiation. Nat Genet 56, 686–696. 10.1038/s41588-024-01678-x

Pongubala, J.M.R., Murre, C., 2021. Spatial Organization of Chromatin: Transcriptional Control of Adaptive Immune Cell Development. Front. Immunol. 12, 633825. 10.3389/fimmu.2021.633825

Posit team, 2024. RStudio: Integrated Development Environment for R. Posit Software, PBC, Boston (MA).

Quinlan, A.R., Hall, I.M., 2010. BEDTools: a flexible suite of utilities for comparing genomic features. Bioinformatics 26, 841–842. 10.1093/bioinformatics/btq033

R Core Team, 2023. R: A Language and Environment for Statistical Computing. R Foundation for Statistical Computing, Vienna, Austria.

Ramírez, F., Ryan, D.P., Grüning, B., Bhardwaj, V., Kilpert, F., Richter, A.S., Heyne, S., Dündar, F., Manke, T., 2016. deepTools2: a next generation web server for deep-sequencing data analysis. Nucleic Acids Res 44, W160–165. 10.1093/nar/gkw257

Ramsköld, D., Wang, E.T., Burge, C.B., Sandberg, R., 2009. An abundance of ubiquitously expressed genes revealed by tissue transcriptome sequence data. PLoS Comput Biol 5, e1000598. 10.1371/journal.pcbi.1000598

Robinson, M.D., McCarthy, D.J., Smyth, G.K., 2010. edgeR: a Bioconductor package for differential expression analysis of digital gene expression data. Bioinformatics 26, 139–140. 10.1093/bioinformatics/btp616

Robinson, M.D., Oshlack, A., 2010. A scaling normalization method for differential expression analysis of RNA-seq data. Genome Biol 11, R25. 10.1186/gb-2010-11-3-r25

Ruiz-Velasco, M., Zaugg, J.B., 2017. Structure meets function: How chromatin organisation conveys functionality. Current Opinion in Systems Biology 1, 129–136. 10.1016/j.coisb.2017.01.003

Sabin, F.R., 1902. On the origin of the lymphatic system from the veins and the development of the lymph hearts and thoracic duct in the pig. Am. J. Anat. 1, 367–389. 10.1002/aja.1000010310

Shannon, P., Markiel, A., Ozier, O., Baliga, N.S., Wang, J.T., Ramage, D., Amin, N., Schwikowski, B., Ideker, T., 2003. Cytoscape: a software environment for integrated models of biomolecular interaction networks. Genome Res 13, 2498–2504. 10.1101/gr.1239303

Shin, H.Y., Smith, M.L., Toy, K.J., Williams, P.M., Bizios, R., Gerritsen, M.E., 2002. VEGF-C mediates cyclic pressure-induced endothelial cell proliferation. Physiological Genomics 11, 245–251. 10.1152/physiolgenomics.00068.2002

Srinivasan, R.S., Dillard, M.E., Lagutin, O.V., Lin, F.-J., Tsai, S., Tsai, M.-J., Samokhvalov, I.M., Oliver, G., 2007. Lineage tracing demonstrates the venous origin of the mammalian lymphatic vasculature. Genes Dev. 21, 2422–2432. 10.1101/gad.1588407

Srinivasan, R.S., Escobedo, N., Yang, Y., Interiano, A., Dillard, M.E., Finkelstein, D., Mukatira, S., Gil, H.J., Nurmi, H., Alitalo, K., Oliver, G., 2014. The Prox1–Vegfr3 feedback loop maintains the identity and the number of lymphatic endothelial cell progenitors. Genes Dev. 28, 2175–2187. 10.1101/gad.216226.113

Srinivasan, R.S., Geng, X., Yang, Y., Wang, Y., Mukatira, S., Studer, M., Porto, M.P.R., Lagutin, O., Oliver, G., 2010. The nuclear hormone receptor Coup-TFII is required for the initiation and early maintenance of *Prox1* expression in lymphatic endothelial cells. Genes Dev. 24, 696–707. 10.1101/gad.1859310

Stanczuk, L., Martinez-Corral, I., Ulvmar, M.H., Zhang, Y., Laviña, B., Fruttiger, M., Adams, R.H., Saur, D., Betsholtz, C., Ortega, S., Alitalo, K., Graupera, M., Mäkinen, T., 2015. cKit Lineage Hemogenic Endothelium-Derived Cells Contribute to Mesenteric Lymphatic Vessels. Cell Reports 10, 1708–1721. 10.1016/j.celrep.2015.02.026

Stuart T, Butler, Hoffman, Hafemeister, Papalexi, Mauck Wm, Hao Y, Stoeckius M, Smibert P, Satija R, 2019. Comprehensive Integration of Single-Cell Data. Cell 177. 10.1016/j.cell.2019.05.031

Tai-Nagara, I., Hasumi, Y., Kusumoto, D., Hasumi, H., Okabe, K., Ando, T., Matsuzaki, F., Itoh, F., Saya, H., Liu, C., Li, W., Mukouyama, Y., Marston Linehan, W., Liu, X., Hirashima, M., Suzuki, Y., Funasaki, S., Satou, Y., Furuya, M., Baba, M., Kubota, Y., 2020. Blood and lymphatic systems are segregated by the FLCN tumor suppressor. Nat Commun 11, 6314. 10.1038/s41467-020-20156-6

van Impel, A., Zhao, Z., Hermkens, D.M.A., Roukens, M.G., Fischer, J.C., Peterson-Maduro, J., Duckers, H., Ober, E.A., Ingham, P.W., Schulte-Merker, S., 2014. Divergence of zebrafish and mouse lymphatic cell fate specification pathways. Development 141, 1228–1238. 10.1242/dev.105031

Vanlandewijck, M., Betsholtz, C., 2018. Single-Cell mRNA Sequencing of the Mouse Brain Vasculature. Methods Mol Biol 1846, 309–324. 10.1007/978-1-4939-8712-2_21

Villefranc, J.A., Amigo, J., Lawson, N.D., 2007. Gateway compatible vectors for analysis of gene function in the zebrafish. Developmental Dynamics 236, 3077–3087. 10.1002/dvdy.21354

Wang, L., Wang, S., Li, W., 2012. RSeQC: quality control of RNA-seq experiments. Bioinformatics 28, 2184–2185. 10.1093/bioinformatics/bts356

Wei, X., Xiang, Y., Peters, D.T., Marius, C., Sun, T., Shan, R., Ou, J., Lin, X., Yue, F., Li, W., Southerland, K.W., Diao, Y., 2022. HiCAR is a robust and sensitive method to analyze open-chromatin-associated genome organization. Mol Cell 82, 1225–1238.e6. 10.1016/j.molcel.2022.01.023

Wigle, J.T., Chowdhury, K., Gruss, P., Oliver, G., 1999. Prox1 function is crucial for mouse lens-fibre elongation. Nat Genet 21, 318–322. 10.1038/6844

Xu, Q., Holder, N., Patient, R., Wilson, S.W., 1994. Spatially regulated expression of three receptor tyrosine kinase genes during gastrulation in the zebrafish. Development 120, 287–299. 10.1242/dev.120.2.287

Yang, H., Luan, Y., Liu, T., Lee, H.J., Fang, L., Wang, Y., Wang, X., Zhang, B., Jin, Q., Ang, K.C., Xing, X., Wang, J., Xu, J., Song, F., Sriranga, I., Khunsriraksakul, C., Salameh, T., Li, D., Choudhary, M.N.K., Topczewski, J., Wang, K., Gerhard, G.S., Hardison, R.C., Wang, T., Cheng, K.C., Yue, F., 2020. A map of cis-regulatory elements and 3D genome structures in zebrafish. Nature 588, 337–343. 10.1038/s41586-020-2962-9

Yaniv, K., Isogai, S., Castranova, D., Dye, L., Hitomi, J., Weinstein, B.M., 2006. Live imaging of lymphatic development in the zebrafish. Nat Med 12, 711–716. 10.1038/nm1427

Yu G (2024). enrichplot: Visualization of Functional Enrichment Result. R package version 1.24.2, [WWW Document], 2024. URL https://yulab-smu.top/biomedical-knowledge-mining-book/. (accessed 8.22.24).

Yu, G., Wang, L.-G., He, Q.-Y., 2015. ChIPseeker: an R/Bioconductor package for ChIP peak annotation, comparison and visualization. Bioinformatics 31, 2382–2383. 10.1093/bioinformatics/btv145

Zanocco-Marani, T., Vignudelli, T., Parenti, S., Gemelli, C., Condorelli, F., Martello, A., Selmi, T., Grande, A., Ferrari, S., 2009. TFE3 transcription factor regulates the expression of MAFB during macrophage differentiation. Experimental Cell Research 315, 1798–1808. 10.1016/j.yexcr.2009.03.018

Zhang, Y., Liu, T., Meyer, C.A., Eeckhoute, J., Johnson, D.S., Bernstein, B.E., Nusbaum, C., Myers, R.M., Brown, M., Li, W., Liu, X.S., 2008. Model-based Analysis of ChIP-Seq (MACS). Genome Biol 9, R137. 10.1186/gb-2008-9-9-r137

